# DiverScan: attention-guided exploratory analysis of animal behaviour across diverse comparative experiments

**DOI:** 10.64898/2026.10.01.755818

**Authors:** Momoe Sukegawa, Hirotsugu Azechi, Tatsumi Butsugan, Nadja Freund, Shiomi Hakataya, Tomohiro Iwata, Noriko Katsu, Koutarou D. Kimura, Hiroki Koda, Akiko Koto, Yuya Makiguchi, Hiroshi Matsui, Koji Miyoshi, Annakarina Mundorf, Hiraku Nishimori, Kazuo Okanoya, Masashi Shiraishi, Susumu Takahashi, Ryoya Tanaka, Genta Toya, Chitose Yamazaki, Shuhei J. Yamazaki, Takuya Maekawa

## Abstract

High-throughput tracking yields rich time-series data of animal behaviour, yet extracting biological insight remains a bottleneck, even for simple control– treatment comparisons. Existing analyses struggle to pinpoint the multiple, nonlinearly intertwined differences hidden in the data and fail to generalise across diverse recording set-ups. Here we present DiverScan, a deep-learning framework for attention-guided exploratory analysis of behavioural time series. DiverScan disentangles condition-dependent behavioural differences by pairing a loss function tailored to data exploration with an attention-branch architecture, and highlights condition-specific behavioural features visually and with explanatory text generated by a large language model. DiverScan is validated on a synthetic benchmark and on datasets from seven laboratories spanning multiple species and both single- and multi-animal behaviour. Beyond recovering known phenotypes, we demonstrate that DiverScan helps researchers generate new hypotheses across these diverse datasets. Available as Jupyter notebooks with a graphical user interface and flexible input formats, DiverScan can integrate into existing behavioural-analysis pipelines.

## 1 Main

High-throughput animal tracking now produces behavioural time series faster than researchers can interpret them. Turning this abundance into biological insight has become a central bottleneck, even for the simple control–treatment comparisons that underpin experimental biology. Researchers typically contrast experimental conditions (e.g., male vs. female, control vs. drug-treated), acquire behavioural data, and analyse them to identify condition-dependent differences. This established pipeline is unchanged, but its data-preparation stage has shifted: animal-tracking tools such as DeepLabCut and the LEAP series[1–4] now yield precise coordinates of animals and their body parts as time series (Fig. 1a). The difficulty of analysing such ‘big behavioural data’ lies in the nature of animal behaviour itself: behaviour is multi-faceted, so the differences between conditions appear as diverse features distributed across individuals, body parts, and time within a high-dimensional, large-scale numerical matrix. If this is overlooked, researchers extract only low-resolution, entangled, or uninterpretable features from otherwise rich data.

**Fig. 1.**
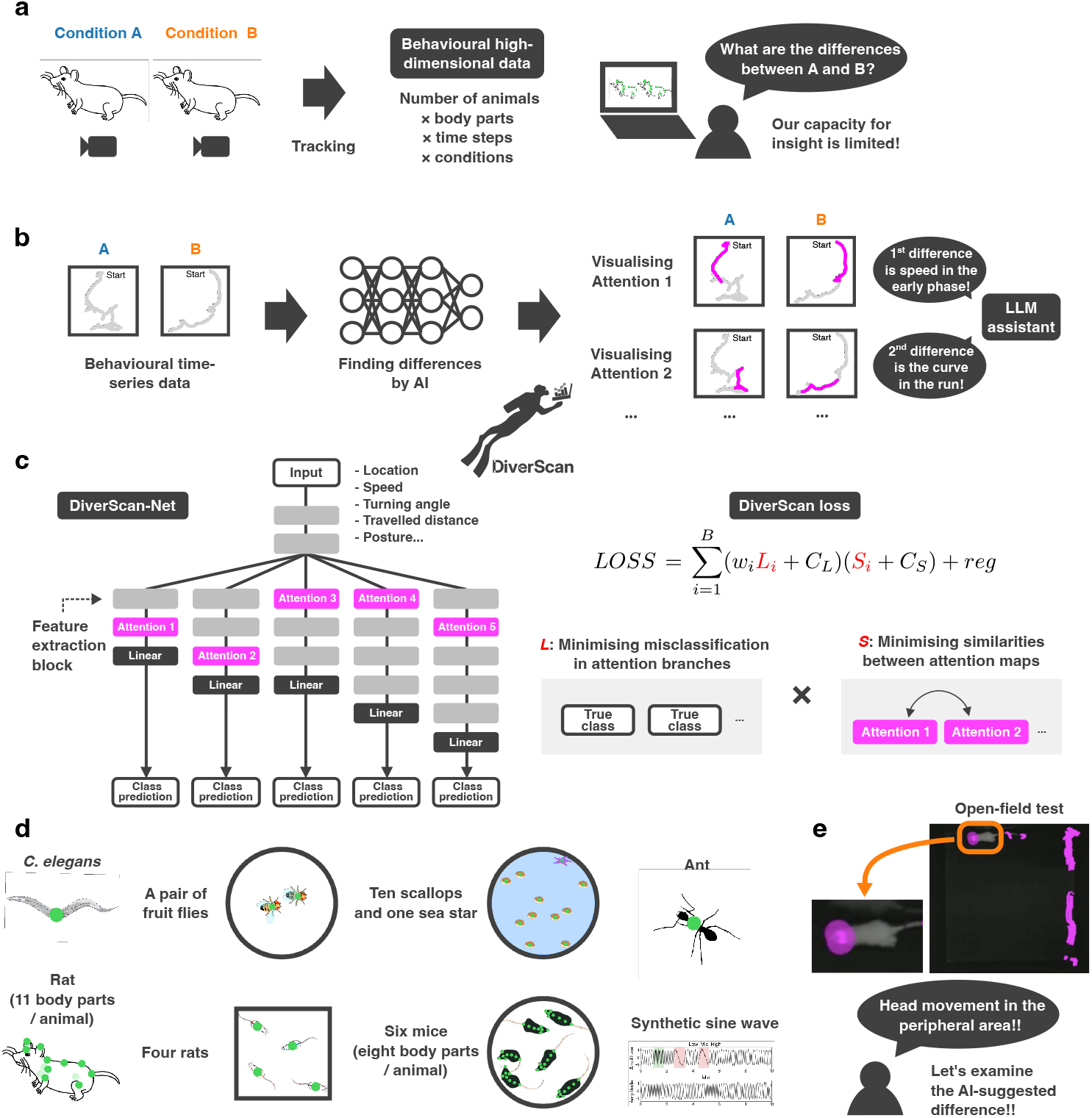
DiverScan, a knowledge-discovery framework based on attention-guided exploratory data analysis. **a**, The challenge addressed by DiverScan. High-throughput tracking now yields high-dimensional behavioural time series across experimental conditions; extracting the condition-dependent differences that matter from this abundance has become a bottleneck. **b,** The DiverScan workflow. Behavioural time series are fed into DiverScan-Net, which learns to discriminate the conditions; the attention of each branch is then visualised separately for every condition (segments with high attention shown in magenta), and an LLM assistant summarises each highlighted difference in natural language (e.g., a speed difference in the early phase, a curvature difference during running). **c,** DiverScan-Net and DiverScan loss. DiverScan-Net comprises *B* parallel branches, each with its own attention layer and linear classifier, together with feature-extraction blocks. DiverScan loss is the product of a classification term *L_i_* (cross-entropy, minimising misclassification within each branch) and a similarity term *S_i_* (minimising the similarity between the branches’ attention maps), so that branches capture diverse, condition-discriminating behaviours. In the single mode, the feature-extraction blocks consist of convolutional or long short-term memory (LSTM) layers; in the multi mode, they consist of spatio-temporal graph convolutional network (ST-GCN) layers. **d,** Applicability of DiverScan across diverse datasets: single *C. elegans* and ants; pairs of fruit flies; groups of four rats; a mixed-species group of ten scallops and one sea star; and skeletal data from a single rat (11 body parts) and six mice (eight body parts), as well as synthetic time series. Green indicates the tracked body parts fed into DiverScan. **e,** An example, drawn from our own analysis of the schizophrenia-model rat (open-field test; see Results), of how DiverScan turns attention into a starting point for exploration. The attention highlights (magenta) point the researcher towards data-driven candidate differences rather than testing only pre-specified metrics.

How, then, should condition-dependent differences be extracted from these large behavioural time series in a rapidly evolving AI landscape? Traditional approaches rely on simple summary statistics or hand-crafted metrics based on expert intuition (Fig. 1a); these are readily interpretable but scale poorly. Machine-learning approaches have begun to address this gap[5–10], yet two difficulties persist. (1) Existing unsupervised approaches (e.g., clustering) yield numerous behavioural motifs but leave how conditions differ to manual follow-up analysis. Supervised approaches (e.g., attention maps[5] or Grad-CAM[11]) directly identify condition-discriminative features but struggle to disentangle the multiple biological differences that coexist in the data. (2) Flexibility is often insufficient for diverse settings: varying numbers of animals, different body-part configurations, or user-defined data beyond pose.

Here we present DiverScan, an attention-guided deep-learning framework that resolves condition-discriminative differences into multiple distinct, spatiotemporally localised behaviours. Given time series from a comparative experiment, DiverScan singles out a set of behavioural features, each capturing a different facet of how the conditions differ, and brings them to the researcher as visual highlights, each with a supporting summary generated by a large language model (LLM) (Fig. 1b). DiverScan operates within one framework whether the target is a single animal (single mode) or multiple animals or body parts (multi mode), and accepts any user-defined time-series data.

The core idea is attention-guided exploratory data analysis (AG-EDA). DiverScan reframes the attention mechanism as a discovery-oriented active compass for exploring big behavioural data, rather than the passive readout used in previous studies. This is realised by an ‘attention-branch’ architecture (DiverScan-Net) and a loss function (DiverScan loss) tailored to data analysis (Fig. 1c). Given a trajectory, DiverScan-Net is trained to predict its experimental condition, so its attention layers learn to attend to condition-discriminative behaviours in a fully data-driven manner. To address point (1), DiverScan-Net has one attention layer per branch, and each attention layer captures a condition-discriminative behaviour that differs from those of the others. To achieve this separation, the structure of each branch is optimised by a neural architecture search, so that the branches apply diverse convolutions to the same input. Critically, DiverScan loss explicitly steers the branches to reduce the similarity of their attentions while maintaining accurate prediction. The attention layers thereby pinpoint the multiple, nonlinearly intertwined differences hidden in the data, each as a separate representation (Fig. 1c). To address point (2), DiverScan loss is defined on the attention representations, allowing the use of various feature extractors suited to the input data. DiverScan thus supports both single- and multi-animal analyses, whether behaviour is represented as single-point trajectories or multi-keypoint poses, and can incorporate user-defined data without changing the framework (Fig. 1d).

We demonstrate DiverScan’s broad applicability using datasets from seven independent laboratories, spanning multiple species and both single- and multi-animal settings: single worms and ants, pairs of fruit flies, groups of four rats, mixed-species groups (ten scallops and one sea star), and skeletal data from a single rat and six mice (Fig. 1d). Across these settings, DiverScan reliably recovered known phenotypes and, going further, helped researchers generate new hypotheses—for example, that mated *Drosophila* females slow down as a virgin male approaches, facilitating the ovipositor-extrusion display by which they signal rejection. Even in previously analysed datasets, DiverScan suggested distinct but unnoticed differences. On a synthetic sine-wave benchmark with known ground truth, it assigned the inserted features to different representations in 100% of correctly classified samples (accuracy *>* 0.96), vs. 2.8% for cross-entropy loss alone (Extended Data Fig. A1 and Supplementary Note). DiverScan is publicly available as Jupyter notebooks on Google Colab with a GUI (Extended Data Fig. A2, https://github.com/M-SUKEGAWA/DiverScan), and accepts DeepLabCut-formatted files as well as user-defined time series (for example, per-timestep behavioural states from other software), integrating into existing pipelines. By turning attention into an engine for exploration, DiverScan surfaces the differences in big behavioural data and shows the researcher where to dive in (Fig. 1e).

## 2 Results

### 2.1 Overview of DiverScan

DiverScan discovers condition-dependent behaviour through supervised classification: given behavioural time series, DiverScan-Net is trained to predict the experimental condition (e.g., control vs. drug-treated), and the attention weights of its attention layers (attention maps) highlight the segments that drive this prediction—when they occur in the single mode, and both which body part or individual and when in the multi mode. These high-attention segments are expected to reflect key behavioural differences between conditions. Each branch of DiverScan-Net has its own attention layer, and the network is trained to minimise DiverScan loss (Fig. 1c), the product of a similarity term and a cross-entropy term. Because both are non-negative, the loss falls only when the branches classify accurately (low cross-entropy) *and* attend to different behaviours (low similarity), so each branch captures a distinct behaviour that discriminates between conditions. The behavioural features identified by DiverScan-Net are then not only visualised but also interpreted in the context of the target species and paradigm. DiverScan supplies an LLM with interpretable cues, the correlations between the attention maps and an exhaustive set of predefined hand-crafted features, and grounds it through retrieval-augmented generation (RAG) over reference materials related to the experiment, yielding biologically grounded, natural-language explanations. These help researchers translate discovered features into concrete, testable hypotheses.

Below, we apply DiverScan to seven datasets from independent laboratories, progressing from single-animal recordings (worms, ants, and rats) to multi-animal and multi-keypoint settings (flies, scallops with a sea star, rats, and mice). Together these span a broad range of comparative designs—two- and three-condition comparisons, attribute-based contrasts such as sex, mixed predator–prey groups, and variable-length recordings—showing that a single framework recovers known findings and surfaces new hypotheses across diverse species, designs, and data formats.

### 2.2 *C. elegans* in an odour-avoidance assay

We first evaluated DiverScan on a well-established paradigm with known signatures, applying its single-mode analysis to the roundworm *C. elegans*. Experience-dependent behavioural changes in worms offer a tractable entry point for studying learning and memory[12]. Using an odour-avoidance assay, we compared two groups of worms during exposure to a repulsive odorant (2-nonanone): worms pre-exposed to the odour (odour-learned) and naive worms with no prior exposure. Previous studies have reported that pre-exposure alters subsequent avoidance behaviour[5, 13–15]: pre-exposed worms keep a greater distance from the odour source, reduce their speed during the relatively early phase, and show lower fluctuations in velocity. We tested whether DiverScan can help researchers both recover these known differences and notice additional, overlooked patterns in each attention branch.

Following the DiverScan workflow (Fig. 2), we prepared centroid trajectories of pre-exposed and naive worms (Fig. 2a; Fig. 3a) and computed behavioural features (e.g., speed, turning angle, distance from the initial location, cumulative travel distance) as single-mode input (Step 1 in Fig. 2b; see Supplementary Table 1 for the input features). We then trained and tested DiverScan-Net with five attention branches (Steps 2– 4 in Fig. 2c–e; see Supplementary Table 1 for the searched hyperparameters) and examined the attention branches through the attention-interpretation module (Step 5 in Fig. 2f). By visualising attention maps and combining them with statistics such as their correlations with hand-crafted features and user-defined attributes, this module offers the researcher both visual attention maps for inspection and natural-language suggestions.

**Fig. 2.**
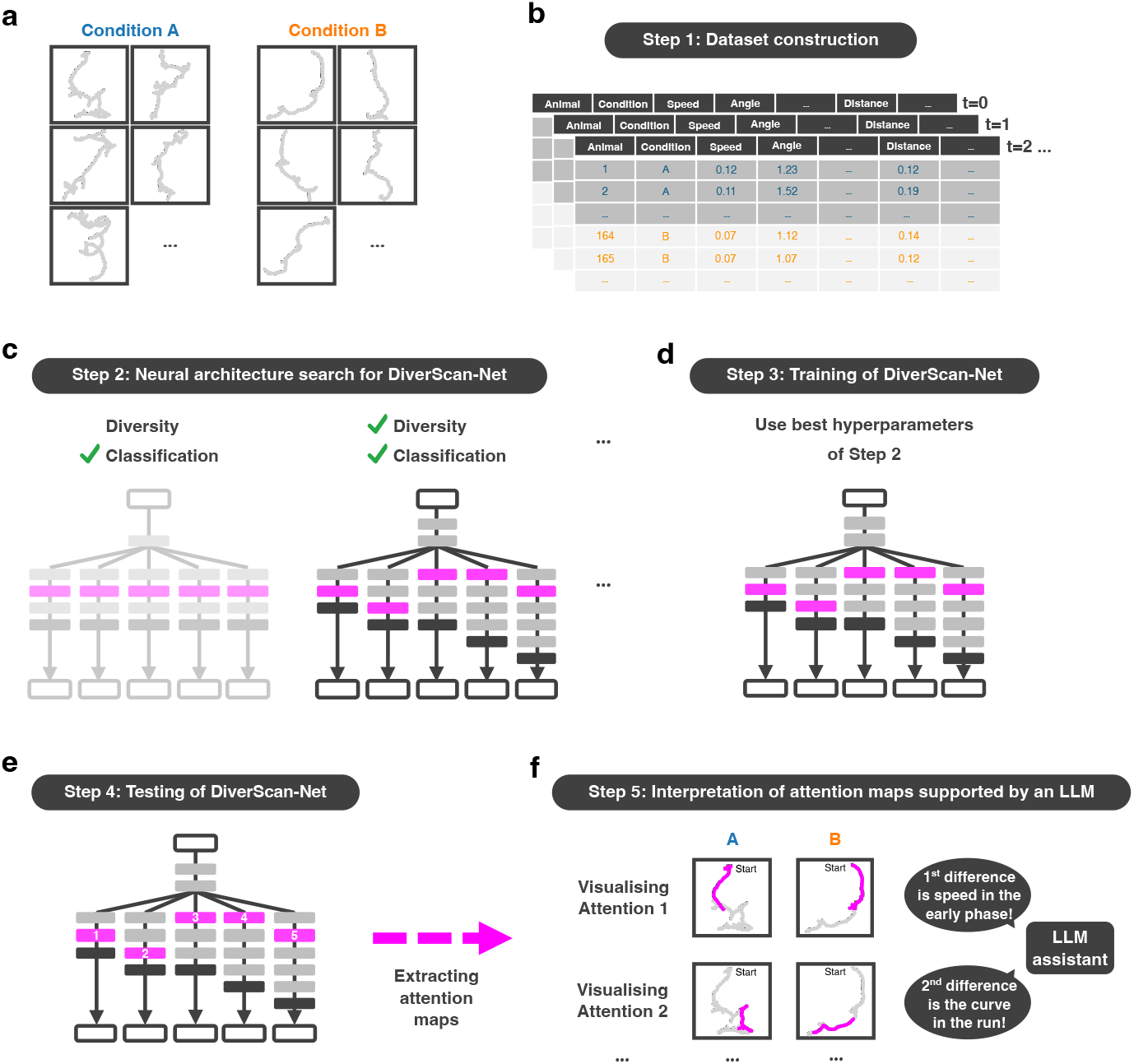
Workflow of DiverScan. **a**, Behavioural time series recorded under two experimental conditions (A vs. B), which serve as the input to the workflow below. **b,** Step 1: Dataset construction. Trajectories are prepared for either the single or the multi mode: in the single mode, each animal’s trajectory is stored as an individual CSV file, whereas in the multi mode, trajectories from multiple animals or body parts are stored together in a single file. DiverScan takes as input not only raw trajectory features such as the *x* and *y* coordinates, but also movement features computed from the trajectories and optional user-defined features (e.g., speed, turning angle, travelled distance, wing angle, and sex). cf. Extended Data Fig. A2a,b. **c,** Step 2: Neural architecture search for DiverScan-Net. DiverScan searches over DiverScan-Net architectures, treating its components as ‘building blocks’. Appropriate hyperparameter settings enable DiverScan-Net to discover diverse features while achieving high classification performance. Alternatively, the user can employ predefined hyperparameters that we have empirically found to perform well. cf. Extended Data Fig. A2c. **d,** Step 3: Training of DiverScan-Net using the best hyperparameters identified in Step 2. cf. Extended Data Fig. A2c. **e,** Step 4: Testing of DiverScan-Net. The network trained in Step 3 is applied to a held-out test set not used in Steps 2 or 3, preventing knowledge discovery from overfitting to the training data. The attention maps for the test set are extracted and stored for Step 5. cf. Extended Data Fig. A2d. **f,** Step 5: Interpretation of the attention maps extracted in Step 4. DiverScan supports this through an attention-interpretation module with two submodules: (i) visual inspection of the attention maps (cf. Extended Data Fig. A2e–h) and (ii) LLM-based assistance via retrieval-augmented generation. See Methods for details.

**Fig. 3.**
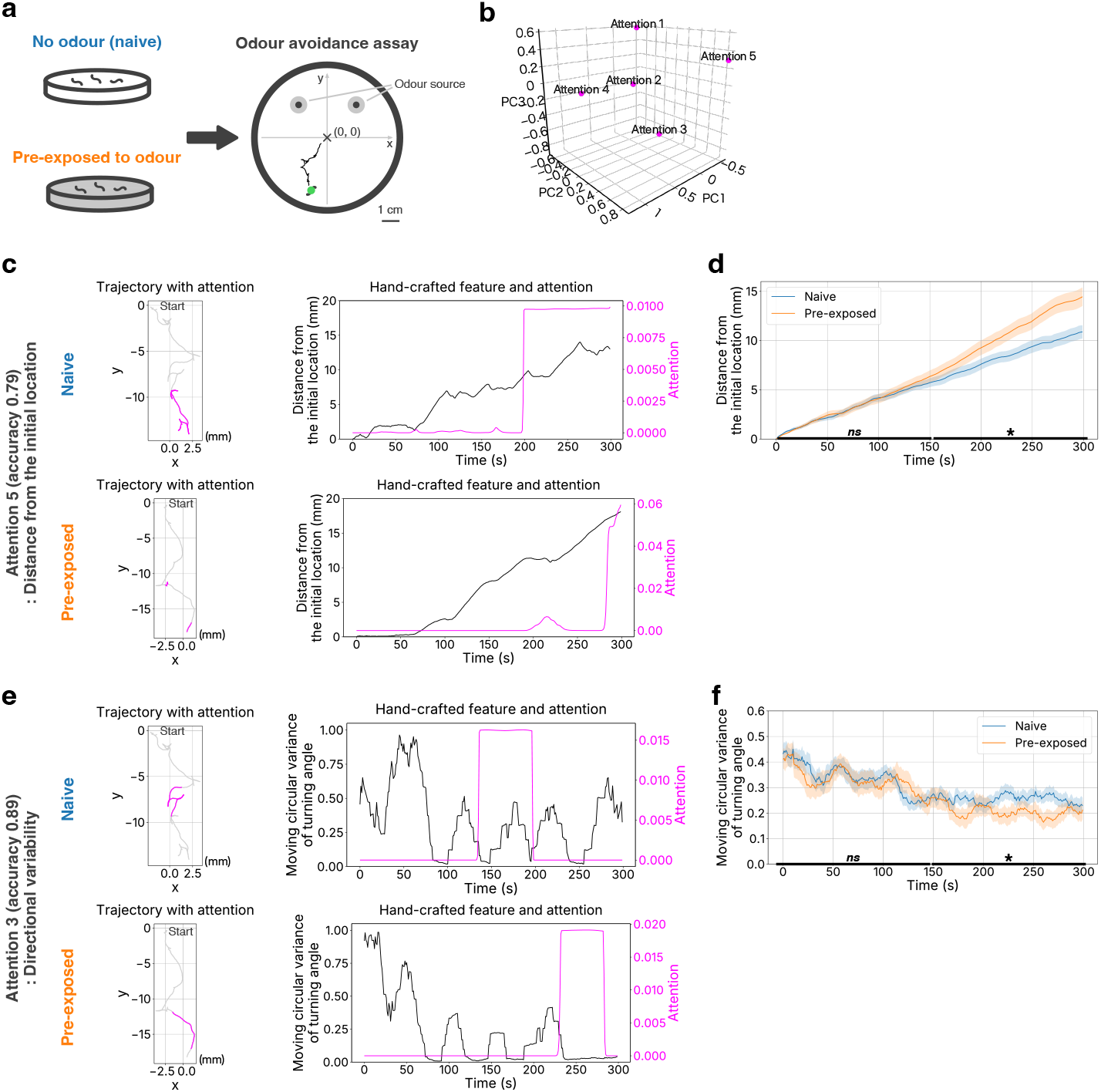
DiverScan recovers a known distance signature and surfaces a turning-variability feature in *C. elegans* odour avoidance. **a**, Odour-avoidance assay comparing naive and pre-exposed worms. The origin (0, 0) is the user-set reference location. **b,** PCA of the five attention branches, summarising their similarities; each point is one branch. **c,** Attention branch 5. Left: example trajectories with high-attention segments in magenta (attention above the chance level 1*/*(*N T ^′^*); *T ^′^* = 300 frames, *N* = 1). Right: time series of the attention value and the distance from the initial location, a hand-crafted feature suggested by DiverScan (moving window, 19 frames). **d,** Distance from the initial location for naive (*N* = 33) and pre-exposed (*N* = 33) worms (mean SEM). Two-sided Student’s *t*-tests with Holm correction on the early (0–150 s) and late (150–300 s) phases (early: adjusted *p* = 0.82, Cohen’s *d* = 0.057, 95% CI [0.91, 0.73]; late: adjusted *p* = 0.043, *d* = 0.58, 95% CI [4.06, 0.33]). Asterisks and *ns* denote *p* 0.05 and *p >* 0.05, respectively. **e,** Attention branch 3, as in **c**. Right: attention value and the moving circular variance of turning angle. **f,** Moving circular variance of turning angle (mean *±* SEM; statistics as in **d**; early: adjusted *p* = 0.71, *d* = 0.093, 95% CI [*−*0.04, 0.06]; late: adjusted *p* = 0.040, *d* = 0.59, 95% CI [0.01, 0.09]).

DiverScan highlighted previously reported differences in four attention branches (1, 2, 4, and 5) and surfaced a difference not emphasised in prior studies (attention branch 3) (Fig. 3, Extended Data Fig. A3). A principal component analysis (PCA) of the attention maps summarised similarities among the attention branches (Fig. 3b; nearby branches capture similar features; see Methods). We first describe the attention branches that recovered well-established differences, then the candidate feature surfaced by attention branch 3.

Attention branch 5 pointed to the best-known signature of this paradigm, the final distance from the odour, highlighting the late phase of the trajectories (Fig. 3c,d). The attention-interpretation module explained:

#### Summary Interpretation of Branch 5

[. . .] For pre-exposed worms, the model pays high attention because they have achieved a large distance from the user-specified location (the odor source [*sic*; the prompt defined this location as the arena centre, near the odour source]). They have actively avoided the smell. [. . .]

Guided by these suggestions of distance-related metrics with late-phase specificity (Supplementary Table 2), our post hoc analysis found that, during the late phase, pre-exposed worms travelled significantly farther from their initial positions than naive worms (Fig. 3d), consistent with acquired odour learning. Curiously, attention branch 5 yielded the lowest classification accuracy of the five attention branches (test accuracy 0.79; see Supplementary Table 3 for per-branch classification performance), suggesting that late-phase distance is comparatively less discriminative within DiverScan.

Attention branch 4 pointed to pre-exposed worms moving more slowly than naive worms during the early phase (Extended Data Fig. A3a,b), consistent with the known short-lived effect of pre-exposure[13]. Attention branches 1 and 2 highlighted reduced velocity fluctuations in pre-exposed worms, through acceleration-based measures (attention branch 1; Extended Data Fig. A3c,d) or variability in travelled distance (attention branch 2; Extended Data Fig. A3e,f); such fluctuations have been reported previously and interpreted as efficient migration[5]. Although earlier work did not specify the relevant time window, DiverScan localised these fluctuations pre-dominantly to the early phase (see Supplementary Table 4 for full statistics), showing that DiverScan can help researchers recover known signatures with high temporal specificity.

Beyond these established differences, attention branch 3 highlighted a fine, potentially overlooked feature. Visual inspection suggested that, within the emphasised segments, pre-exposed worms followed smoother, more gradually curving paths than naive worms (Fig. 3e). Consistent with this, the module explained:

#### Summary Interpretation of Branch 3

[. . .] Attention 3 tells a distinct behavioral story: to identify a pre-exposed worm, the model waits until the later stages of the experiment (Time) and looks for worms that have traveled a long distance with smooth, non-erratic movements (Low Turn Variance). [. . .]

Building on this suggestion of turning-related metrics with late-phase specificity (Supplementary Table 2), our post hoc analysis showed that pre-exposed worms had reduced moving circular variance of turning angle during the late phase, indicating smaller moment-to-moment directional changes (Fig. 3f). This raises the possibility that pre-exposed worms stabilise their avoidance strategy by the late phase, perhaps reflecting a more settled internal state—an increased ‘confidence’—whereas naive worms remain exploratory. It motivates a testable hypothesis: that learning stabilises decision-making by reducing behavioural variability. This candidate had not been identified in extensive previous work, suggesting that DiverScan can point to new hypotheses even in a well-characterised paradigm. Notably, this attention branch achieved the highest classification accuracy (0.89) of the five. That the best-known signature (attention branch 5) was the least discriminative, whereas this overlooked feature was the most, underscores the limitations of hand-crafted metrics based on human intuition.

Overall, DiverScan helped us recover and surface multiple signatures of experience-dependent odour avoidance in *C. elegans* with clear temporal localisation, supporting researchers in exploring diverse behavioural features through its attention-guided exploratory mechanism.

### 2.3 Courtship behaviour in fruit flies

Next, we evaluated DiverScan’s multi mode for analysing group-level dynamics. Studying group behaviour is challenging because it requires accounting for interactions among individuals. DiverScan’s multi mode addresses this by operating on time-series graphs: nodes represent individuals characterised by behavioural-state features and animal-specific attributes (e.g., position, speed, pose, and sex), and edges represent pairwise relations (e.g., inter-animal distance) that evolve as animals move. Operating directly on these spatiotemporal features via the attention mechanism, DiverScan supports fine-grained analysis of group dynamics.

To evaluate the multi mode, we began with a two-agent interaction: courtship behaviour in two fruit flies, *D. melanogaster*. Fruit-fly courtship is a tractable model for investigating how reproductive behaviours are programmed in the nervous system[16]; the male follows the female and performs displays such as wing extension and courtship song. Previous studies have shown that virgin males prefer virgin females over mated females and court them more strongly (e.g., increased following and wing extension towards virgin females)[17], and that mated females are more resistant to remating (e.g., moving away, ovipositor extrusion against males)[18]. We prepared two conditions: pairs of a virgin female and a virgin male, and pairs of a mated female and a virgin male (Fig. 4a). DiverScan automatically constructed time-series graphs, representing each animal as a node and their relation (e.g., distance between two flies) as an edge connecting the two nodes (Fig. 4a, green lines and dots). Each node carried the fly’s apparent wing angle (estimated by FlyTracker[19], defined as the maximum of the left and right wing angles relative to the body axis) as a pose feature.

**Fig. 4.**
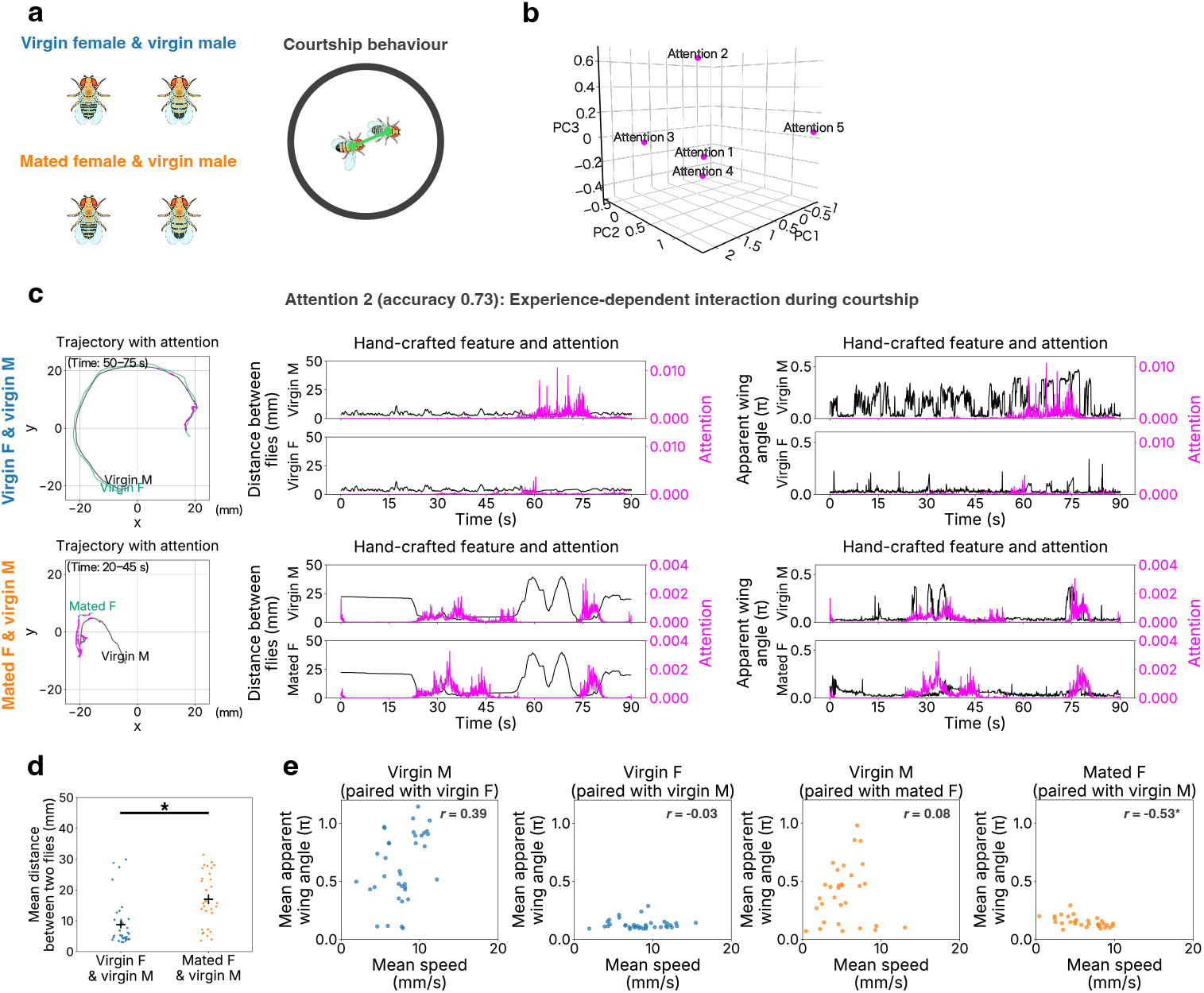
DiverScan surfaces an experience-dependent rejection behaviour in fruit-fly courtship. **a**, Schematic of the two conditions: a virgin female with a virgin male, and a mated female with a virgin male. DiverScan automatically constructed time-series graphs, representing each fly as a node and their relation (e.g., distance between two flies) as an edge connecting the two nodes. **b,** PCA of the five attention branches; each point is one attention branch. **c,** Attention branch 2. Left: example trajectories with high-attention segments highlighted in magenta (attention above the chance level 1*/*(*N T ^′^*); *T ^′^* = 2,700 frames, *N* = 2 animals). Middle and right: time series of the attention value together with the inter-fly distance (middle) and the apparent wing angle (right), for the male (M) and female (F). **d,** Mean inter-fly distance for virgin-female–virgin-male (*N* = 36 clips) and mated-female–virgin-male (*N* = 31 clips) pairs. Here, a clip is a segment of a recording, used to enable deep-learning-based exploratory analysis on a limited number of recordings (see Discussion and Methods for its effect on the analysis). Horizontal and vertical bars indicate the mean and SEM, respectively. Two-sided Welch’s *t*-test: *p* = 1.1 10*^−^*^4^, Cohen’s *d* = 1.0, 95% CI [12.19, 4.26]. Asterisks denote *p* 0.05. **e,** Mean apparent wing angle vs. mean speed for pairs in close proximity (inter-animal distance 17.0 mm; see **d**), with the Pearson correlation *r* and Holm-corrected *p* for each group: virgin M paired with virgin F (*r* = 0.39, adjusted *p* = 0.058, 95% CI [0.07, 0.64]), virgin F paired with virgin M (*r* = *−*0.03, adjusted *p* = 1, 95% CI [*−*0.36, 0.30]), virgin M paired with mated F (*r* = 0.08, adjusted *p* = 1, 95% CI [*−*0.28, 0.43]), and mated F paired with virgin M (*r* = *−*0.53, adjusted *p* = 0.0086, 95% CI [*−*0.74, *−*0.22]). The 95% CIs are not adjusted for multiple comparisons. Asterisks denote adjusted *p ≤* 0.05.

We present the results in Fig. 4 and Extended Data Fig. A4. DiverScan helped us (i) surface an unexpected fine-grained social behaviour and (ii) recover previously known features across its attention branches (overview in Fig. 4b).

We first describe attention branch 2, which captured a characteristic interaction pattern in mated females, represented by a combination of basic behavioural metrics. Based on the attention maps and their correlations with hand-crafted features (Fig. 4c, Supplementary Table 5), the attention-interpretation module explained:

- **VM–MF [pairs of a mated female and a virgin male]:** Highly correlated with max wing ang (up to 0.208) and strongly negatively correlated with Distance between nodes (−0.301) and Speed (−0.104). This indicates the attention is sharply focused on moments when the male is very close to the female, moving slowly, and extending his wing [. . .].
- **VM–VF [pairs of a virgin female and a virgin male]:** The strong wing angle correlation disappears. Instead, attention is focused on negative Distance between nodes (−0.100) and negative Speed/Acceleration.

The module noted that, although these correlations ranked relatively highly among the candidate features, their magnitudes were modest, suggesting that the effect may arise from combinations of features and/or nonlinear interactions. Guided by these suggestions, we hypothesised that attention branch 2 highlights a relationship between wing posture and speed when the animals are close together, and performed a post hoc analysis focusing on periods of close proximity. Consistent with previous studies, mated-female and virgin-male pairs maintained greater inter-animal distances, indicative of reduced courtship (Fig. 4d). Using a distance threshold defined as the mean inter-animal distance for these pairs, we found that when the virgin male was nearby, mated females tended to slow down as the apparent wing angle increased (Fig. 4e). This appeared inconsistent with prior reports that mated females typically move away from males[18], so we manually inspected the recorded videos. The inspection revealed that the increase in apparent wing angle did not reflect active wing opening; rather, because the apparent wing angle is defined relative to the body axis, it resulted from a shift in the estimated body axis during ovipositor extrusion towards the male. In *D. melanogaster*, ovipositor extrusion in mated females is a well-described rejection behaviour elicited by male approach[20], whose cues could serve as a rejection display[17]. Together, these observations motivate a new hypothesis: that mated females slow down to facilitate rejection signalling during male approach.

The remaining attention branches pointed to more conventional features. Consistent with prior reports[17], attention branch 3 indicated that virgin males walked faster when paired with virgin females (i.e., stronger courtship) than with mated females (Extended Data Fig. A4a,b). On the female side, virgin females also showed a higher mean speed than mated females (Extended Data Fig. A4b); one explanation is that virgin females rarely perform ovipositor extrusion, so males continue to chase them, raising the mean female walking speed. Reflecting this enhanced courtship, attention branch 4 highlighted the periodicity of the pair’s locations in the round arena: virgin-male and virgin-female pairs moved circularly, possibly along the perimeter (Extended Data Fig. A4c,d). Attention branch 1 resembled attention branch 4, as expected from the PCA result (Fig. 4b), representing this circular motion through angle-related features (Supplementary Table 5). Finally, attention branch 5 focused on apparent wing angle, particularly its accumulation on the female side (Extended Data Fig. A4e,f); here, however, classification accuracy was 0.61 (Supplementary Table 3) and the comparison between virgin and mated females was modest (adjusted *p* = 0.18, Cohen’s *d* = 0.33, 95% CI [−0.03, 0.01]; Supplementary Table 4; Extended Data Fig. A4f). Since virgin females rarely exhibit ovipositor extrusion whereas mated females do so more frequently[17], this may underlie the mating-status-dependent difference in apparent wing angle; within DiverScan, classification accuracy likely reflects, at least in part, the strength of the underlying difference. Together, these attention branches are consistent with previous studies, confirming that DiverScan operates on multi-animal data.

Overall, DiverScan recovered known single-animal features and surfaced a complex, interaction-level feature in fruit-fly courtship, supporting the exploration of multi-animal dynamics and new hypotheses about social interaction.

### 2.4 Sex-dependent predator-avoidance strategies in scallops

To probe DiverScan’s flexibility, we applied its multi mode to predator–prey data requiring both a multi-group comparison and user-defined node features. The Japanese scallop (*Mizuhopecten yessoensis*) is an ecologically and commercially important bivalve whose populations are strongly influenced by sea star predation, so understanding its anti-predator strategies can inform both ecological theory and aquaculture management. As filter feeders, scallops typically move only short distances and adjust their spacing on the seafloor; when threatened, however, they show a distinctive escape response, shifting to rapid swimming jumps. They also occur at high densities (e.g., three per m^2^ in natural habitats[21]), suggesting that interactions among neighbours may influence predator avoidance and survival. Yet predator-avoidance strategies in scallop groups remain poorly understood. To address this, we used DiverScan’s multi mode to investigate predator avoidance in scallop groups, focusing on sex differences

(Fig. 5a). Japanese scallops are protandrous hermaphrodites—first maturing as males, then transitioning to females—and maintain an approximately 1:1 sex ratio, so sex-dependent strategies under predation, if present, would offer important insights into bivalve ecology. We prepared three conditions, each with one sea star: ten males, ten females, and five males with five females (Fig. 5a). Species identity (scallop vs. sea star) was encoded as a binary node feature.

**Fig. 5.**
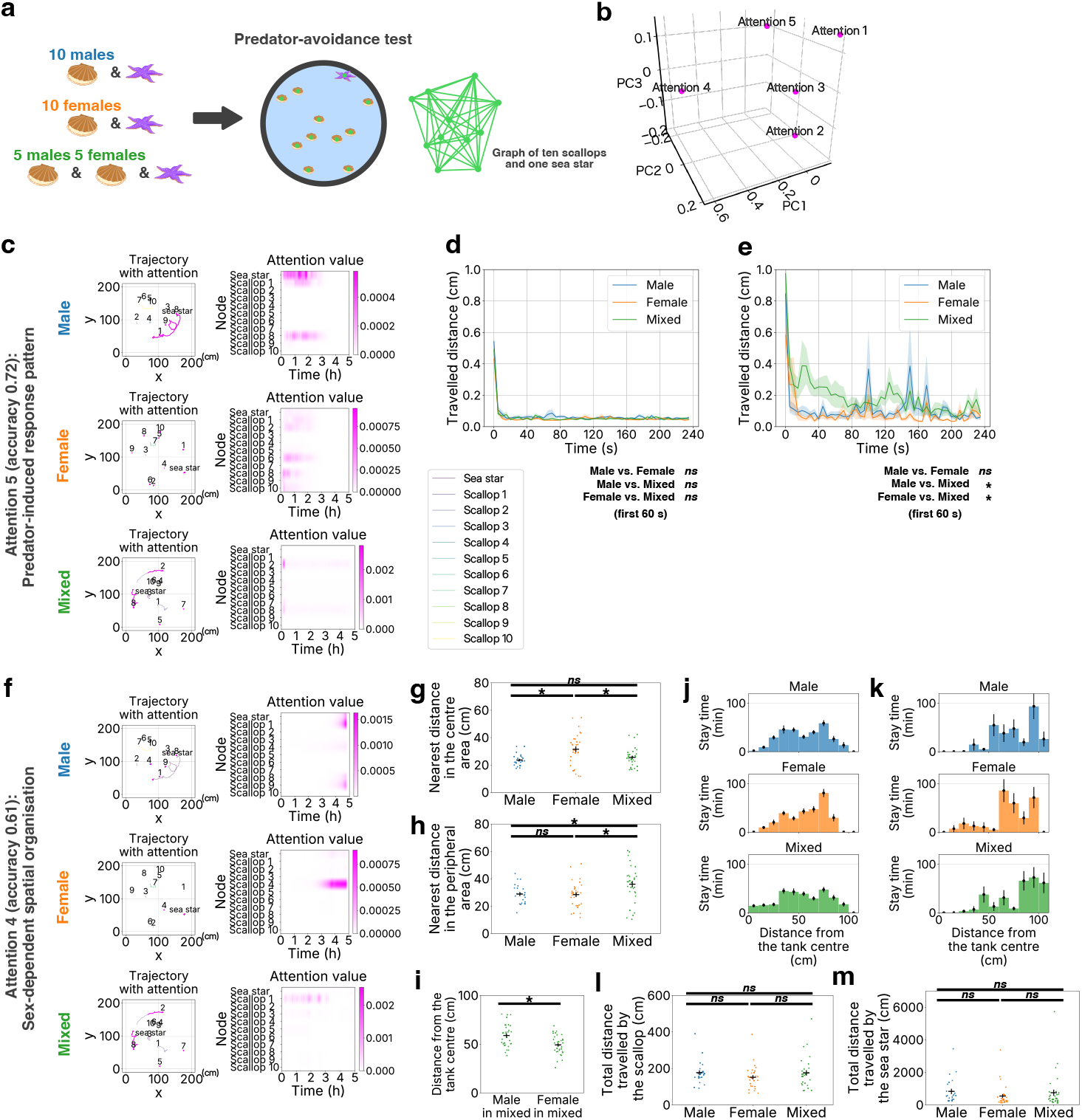
DiverScan reveals sex-dependent predator-avoidance strategies in scallops. Throughout, attention above the chance level 1*/*(*N T ^′^*) is shown in magenta, where *T ^′^* is the number of valid time steps (*T ^′^* = *T* = 3,600 frames here) and *N* = 11 nodes (ten scallops and one sea star). Statistical details are given in Supplementary Table 4; two-sided *t*-tests with Holm correction were used, and asterisks and *ns* denote *p* 0.05 and *p >* 0.05, respectively. Horizontal and vertical bars (or solid lines and shaded regions) indicate the mean and SEM, respectively. **a,** Predator-avoidance test; scallops and a sea star are nodes of the time-series graph. **b,** PCA of the five attention branches. **c,** Attention branch 5. Left: example trajectories with high-attention segments; right: heatmaps of attention values. **d,** Scallop movement in the absence of sea-star movement (male, *N* = 22 clips; female, *N* = 30; mixed, *N* = 32). **e,** Scallop movement after predator-movement onset (male, *N* = 8 clips; female, *N* = 6; mixed, *N* = 8; only clips containing at least one period of sea-star movement were analysed). **f,** Attention branch 4. Left: example trajectories with high-attention segments; right: heatmaps of attention values. **g,** Nearest-neighbour distance between scallops in the central area (within 50 cm of the centre of the tank, which has a 97 cm radius), measured to the closest conspecific in either area (male, 23 clips; female, 30; mixed, 32). **h,** Nearest-neighbour distance between scallops in the peripheral area (beyond 50 cm from the centre). **i,** Mean distance from the tank centre in the mixed condition, comparing male (*N* = 32 clips) and female (*N* = 32 clips) scallops. **j,** Spatial distribution of scallops (per scallop). **k,** Spatial distribution of sea stars (per sea star). **l,** Total distance travelled by scallops (per scallop). **m,** Total distance travelled by sea stars (per sea star).

We describe representative attention branches 4 and 5: attention branch 5 attended to active predator responses, whereas attention branch 4 attended to static spatial-distribution patterns shaped by the group’s sex composition (cf. Fig. 5b). For the post hoc analyses, scallops preyed upon during the experiments (*N* = 15) were excluded to avoid biases (5 males, 5 females, and 4 males with 1 female in the male, female, and mixed conditions, respectively).

Attention branch 5 highlighted sea-star activity and scallops located near sea stars in its attention maps (Fig. 5c, Supplementary Table 6). Prompted by this, we compared scallop movement between periods when the sea star was active and inactive (defined by a speed threshold; see Methods), measuring the mean displacement per scallop. During periods without sea-star movement, scallops in all conditions stopped moving almost immediately after an initial movement (Fig. 5d). By contrast, during periods of sea-star movement the scallops kept moving for longer; in particular, scallops in the mixed condition sustained active movement responses to sea-star activity more than those in the male-only and female-only conditions, especially during the first 60 s after predator movement began (male vs. mixed, adjusted *p* = 0.0092, Cohen’s *d* = 1.8, 95% CI [−0.30, −0.08]; female vs. mixed, adjusted *p* = 0.018, *d* = 1.6, 95% CI [−0.35, −0.06]; Fig. 5e, Supplementary Table 4). This raises the possibility of a sex-dependent avoidance strategy in scallops, an under-studied area of behavioural research in bivalves. We discuss this further with attention branch 4.

Attention branch 4 appeared to capture the spatial organisation of scallops within the tank (Fig. 5f). DiverScan’s attention-interpretation module suggested the distance from the tank centre and the mean inter-individual distance as the primary features driving this attention (Supplementary Table 6), indicating that scallop spacing may vary with position in the tank. Post hoc analyses suggested that scallops maintained greater spacing in the central region in the female condition, and peripherally in the mixed condition (Fig. 5g,h). In the mixed condition, further analysis pointed to sex-based spatial segregation, with males more likely than females to occupy peripheral areas (*p* = 8.5 × 10^−4^, Cohen’s *d* = 0.88, 95% CI [4.12, 15.04]; Fig. 5i, Supplementary Table 4). Overall spatial occupancy and total distance travelled did not differ across conditions (Fig. 5j–m, Supplementary Table 4), suggesting that these patterns are unlikely to reflect basic locomotor activity.

Taken together, the heightened predator response and spatial segregation in the mixed condition point to a possible link between group composition and collective defensive capacity, which we interpret cautiously in light of several possibilities. First, the spatial patterns may reflect inherent behavioural differences between the sexes: comparable sex-specific preferences have been reported in other bivalves, with females occupying shallower depths than males[22], and under high density, spatial optimisation matters for filter feeders competing for suspended food[23]. Because the experiments fell in the non-reproductive period, direct effects of mating are unlikely[24]. Second, the adaptive value of the enhanced reactivity is uncertain, given the vigilance–energy trade-offs documented in gastropod anti-predator behaviour[25]: greater reactivity need not improve survival if energetically costly. Third, because sex composition can shape group structure and defence in other marine invertebrates[26], males’ tendency to occupy peripheral positions may reflect the coexistence of sex-specific strategies rather than altruism. Finally, we cannot exclude that tank design and drainage structures differentially affected the sexes’ spatial preferences.

Despite these alternative interpretations, marked sex differences in predator response and space use were observed consistently across two behavioural metrics, pointing to the hypothesis that scallops adopt sex-specific adaptive strategies and that sex composition helps shape group-level dynamics in marine bivalves. DiverScan surfaced them from a group of eleven non-model animals that included a predator, demonstrating that its exploratory mechanism generalises to non-model groups under ecologically complex conditions.

### 2.5 Application of DiverScan in various experimental settings

Furthermore, we validated DiverScan across a broader variety of experimental settings. **Activity of isolated ants.** We next applied DiverScan’s single mode to ants subjected to social isolation[27, 28], a severe stressor for social animals. Here the trajectory length varied with the duration of ‘trips’ (an ant leaving the nest and returning) and with tracking gaps; DiverScan accommodates such data because it processes variable-length time series. We compared socially isolated ants with and without melatonin treatment, which mitigates isolation-induced stress (control and melatonin-treated, respectively; Fig. 6a). The attention maps (Fig. 6b) were negatively correlated with azimuth (i.e., direction of movement; Fig. 6c, Extended Data Fig. A5a) and, relatedly, with the distance from the nest entrance (Extended Data Fig. A5a, Supplementary Table 7). Building on these suggestions and our prior knowledge that the food and water sources were placed far from the nest entrance, at negative *x* coordinates, we observed that the azimuth distribution differed between treatments in the region *x <* −20: control ants preferentially moved towards the food source, whereas melatonin-treated ants were biased towards the water source (Fig. 6d). Although conventional metrics based on time spent in the food and water areas revealed no difference between treatments, DiverScan pointed to a spatial bias towards specific landmarks in isolated individuals. See Supplementary Note and Extended Data Fig. A5 for detailed discussion.

**Fig. 6.**
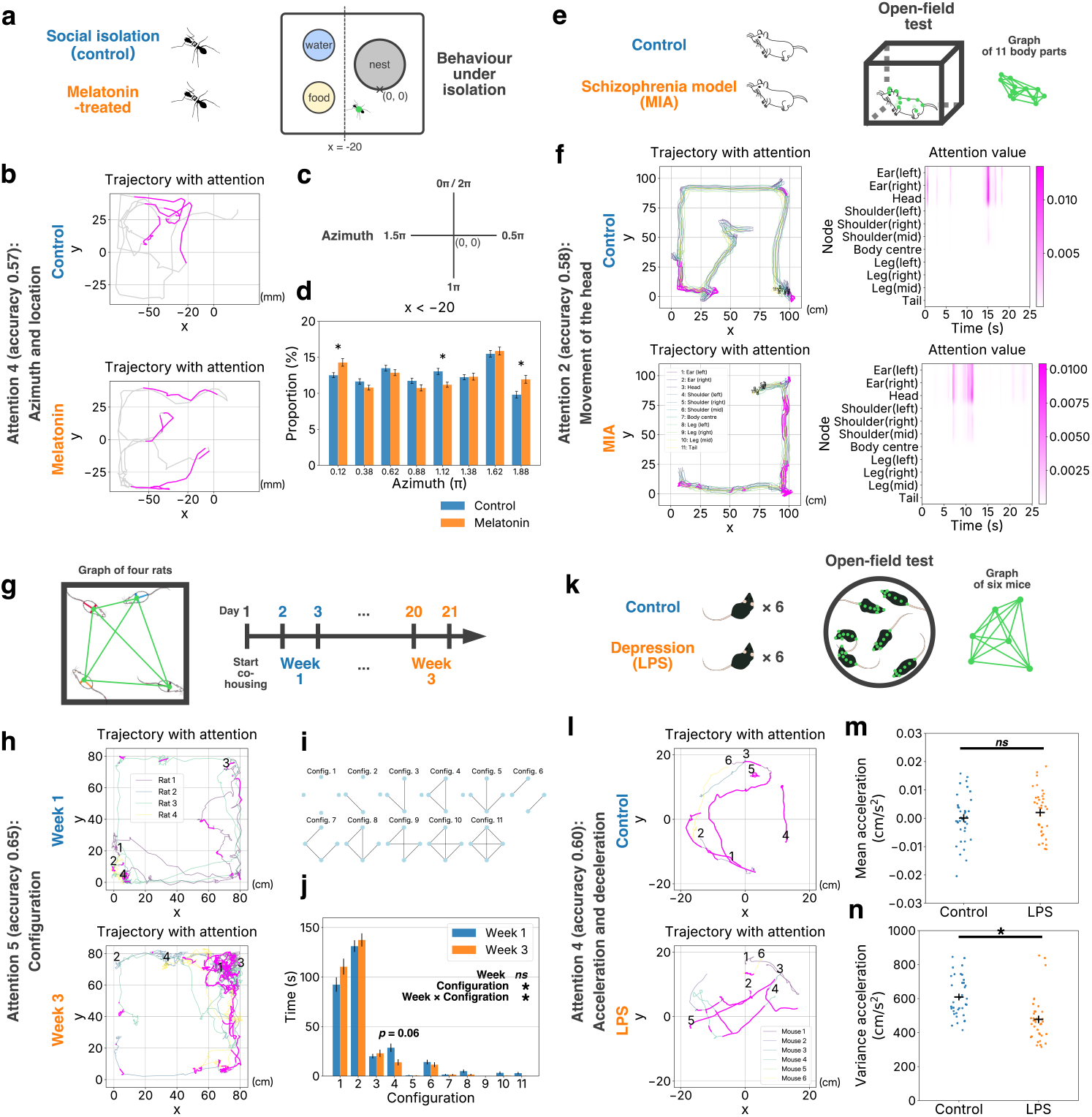
DiverScan generalises across diverse species, input formats, and comparative designs. Throughout, attention above the chance level 1*/*(*N T ^′^*) is shown in magenta, where *T ^′^* is the number of valid time steps; statistical details are given in Supplementary Table 4, and asterisks and *ns* denote *p* 0.05 and *p >* 0.05, respectively. Solid lines and shaded regions (or horizontal and vertical bars) indicate the mean and SEM, respectively. **a,** Social isolation in ants, with or without melatonin. Ants moved freely in a box with the nest entrance at the origin (0, 0) and the food and water sources on the left. **b,** Attention branch 4: example trajectories with high-attention segments (upper, *T ^′^* = 228; lower, *T ^′^* = 195; 0.5 s per frame). The line *x* = 20 separates the food-and-water area from the nest area for the analysis. **c,** Definition of azimuth (direction of movement). **d,** Proportion of azimuths in the region *x <* 20 (445 control vs. 506 melatonin-treated sub-trips). **e,** Open-field test in a rat model of schizophrenia (11 body parts per rat); body parts are nodes of the time-series graph. **f,** Attention branch 2: example trajectories (left) and heatmaps of attention values (right). **g,** Three-week co-housing of four rats; each rat is a node of the time-series graph. **h,** Attention branch 5: example trajectories with high-attention segments. **i,** Eleven configurations of four rats. Lines denote pairs that are ‘near’ (inter-rat distance 15 cm). **j,** Time spent across the eleven configurations (week 1, *N* = 112 clips; week 3, *N* = 112 clips). **k,** Open-field test in a mouse model of depression (eight body parts per animal: nose, left ear, right ear, head, neck, body centre, body rear, and tail base); animals are nodes of the time-series graph. **l,** Attention branch 4: example trajectories with high-attention segments (10 s windows from300 s clips). **m,** Mean acceleration (control, *N* = 36 clips; depression, *N* = 36). **n,** Variance of acceleration (*N* as in **m**).

#### Behaviour of a rat model of schizophrenia

We also applied DiverScan’s multi mode to a tracking set-up capturing multiple body parts of a single animal, now easily collected by recent tracking software. In the multi mode, DiverScan treats these as a skeleton, building time-series graphs in which nodes are body parts and edges are inter-part distances. We tested this on a rat model of schizophrenia (Fig. 6e), tracking 11 body parts per rat. Using DiverScan, we noted that fine, short-timescale head-tilting motions appeared characteristic of the schizophrenia-model rats, because attention tended to fall on the rats’ head-related body parts (Fig. 6f, Extended Data Fig. A6, Supplementary Table 8). This raises the possibility of impaired motor coordination, a key but often overlooked symptom of schizophrenia[29] not detected in previous analyses[30]. By surfacing fine head movements that earlier analyses had missed, DiverScan supports the broader use of multi-body-part tracking datasets. See Supplementary Note and Extended Data Fig. A6 for further discussion.

#### Social relationship formation in rats

We next applied DiverScan’s multi mode to a group of four rats. Rats form complex social relationships[31, 32] thought to develop through formation and maintenance phases, with encounters among unfamiliar individuals destabilising interaction patterns early on[33, 34]. Representing each rat as a node, we compared an early phase (week 1) with a late phase (week 3) of co-housing (Fig. 6g), restricting the analysis to periods when the rats were dynamically active (see Methods). For attention branch 5, the attention-interpretation module pointed to inter-rat distances and the spatial-configuration pattern of the four rats (Fig. 6h, Supplementary Table 9); guided by this, we examined the 11 possible spatial configurations of the group (Fig. 6i) and found a week-by-configuration interaction (Fig. 6j): configuration 4—three rats together with one apart—tended to last longer in week 1, with the same individual consistently separated (Extended Data Fig. A7a). Consistent with Weiss et al.[35], this ‘three-together, one-solo’ pattern may typify early relationship formation. Simple metrics such as the mean inter-rat, nearest-neighbour, and farthest-neighbour distances did not differ between weeks (Extended Data Fig. A7b–d), yet DiverScan pointed to group-level spatial patterns that these metrics missed.

#### Behaviour of a mouse model of depression

Finally, we applied DiverScan’s multi mode to multi-animal, multi-keypoint tracking; here DiverScan-Net converts body-part coordinates into pose features and uses them as node features. We applied this to an open-field test with six mice in a mouse model of depression (Fig. 6k). Guided by the attention-interpretation module, which suggested speed, acceleration, and their variance, we observed that, relative to controls, the depression-model mice tended to show lower acceleration variability and lower mean speed, whereas mean acceleration did not differ between conditions (Fig. 6l–n, Extended Data Fig. A8a,b, Supplementary Table 10). The depression-model mice also appeared to transition into and out of social isolation less frequently, remaining in longer isolation bouts even though their total time isolated was unchanged (Extended Data Fig. A8c,d). These observations motivate a hypothesis that, beyond the well-known immobility phenotype, the abruptness of movement may serve as a depression-like indicator, potentially reflecting psychomotor retardation and behavioural flattening[36]. DiverScan performed well here, although it did not surface strong inter-animal or posture-related features—likely because the arena (40 cm in diameter, six mice) offered limited space for interaction.

## 3 Discussion

Analysis and interpretation of large-scale behavioural time series remain major challenges in behavioural science. Here we developed DiverScan, a framework for interpretable and flexible exploration of behavioural differences in comparative designs. Across diverse settings—single-animal trajectories, multi-animal interactions, user-defined attributes and time series, and synthetic data—DiverScan consistently supported meaningful knowledge discovery.

A key conceptual contribution is attention-guided exploratory data analysis (AG-EDA) for animal behavioural time series. Whereas many existing attention-based methods are optimised primarily for prediction, DiverScan explicitly shapes exploration through its loss function, driving the network to separate diverse, behaviourally meaningful features. Although several explainable-AI (XAI) approaches visualise multiple attentions or employ methods such as Grad-CAM[11], they typically rely on standard objectives and architectures, treating attention as a post-hoc readout. Such approaches face the limitation summarised as ‘attention is not explanation’[37]: attention weights do not necessarily yield useful explanations. In contrast, DiverScan reframes attention as an actively optimised analytical probe rather than a passive by-product. It steers this probe through both the loss and the architecture under exploration-oriented constraints, separating the meaningful, distinct representations in the data. In DiverScan, attentions are starting points that the user interrogates— in dialogue with a RAG-grounded LLM, via the attention maps, through post hoc analysis, and, where needed, by direct inspection of the recordings—as in our fruit-fly analysis, where an apparent wing-angle signal was traced to ovipositor extrusion only after re-examining the videos. Together, these components turn attention into an instrument for pinpointing the differences between conditions, letting users examine a dataset from multiple perspectives and form new hypotheses grounded in both data-driven signals and biological knowledge.

DiverScan complements, rather than replaces, existing behavioural-analysis tools: its flexible input formats make it a natural partner for tools that perform action segmentation and pose representation[6, 9, 38, 39]. Within an exploratory pipeline, DiverScan sits at a later stage—the analysis step itself: once such tools have summarised behavioural data, DiverScan can analyse those summaries alongside additional modalities. We therefore see DiverScan as one component of a broader, collaborative ecosystem of behavioural-analysis methods.

Several limitations should be noted. First, the regularisation that finds distinct, discriminative features can trade off against computational cost, requiring careful hyperparameter tuning; we provide predefined hyperparameters, but reducing this cost is left to future work. Second, what counts as a ‘difference’ is context-dependent, and task-appropriate similarity metrics may need to be chosen for the scientific question. Third, as a deep-learning method, DiverScan is data-hungry: classical analyses can, under appropriate assumptions, be applied to small samples, whereas DiverScan requires larger datasets, and with limited data the risk of false positives rises. The features DiverScan surfaces are therefore candidates for investigation, not confirmed findings: each should be examined further—through post hoc analysis, independent data, or follow-up experiments—before being accepted as genuine. We therefore position DiverScan as a tool for exploratory rather than confirmatory analysis.

In summary, DiverScan endows attention-based deep learning with an attention-guided strategy and integrated interpretability for exploratory data analysis. By treating attention as an instrument rather than a readout, DiverScan offers a practical bridge between exploration and explanation, with broad potential across behavioural science.

## 4 Methods

The DiverScan code, installation instructions, and demos are available as described in the Code availability section; the entire workflow runs on Google Colab without local installation. We welcome community comments and contributions through the GitHub repository.

### 4.1 Data preparation

#### 4.1.1 *C. elegans* in an odour-avoidance assay

The odour-avoidance assay in *C. elegans* was performed as previously described[14], using the same dataset as in Maekawa et al.[5]. Ethical approval was not required for this data collection. Worms avoid the repulsive odorant 2-nonanone, a volatile compound reported to be produced by *P. aeruginosa*, a bacterial pathogen of *C. elegans*[40]. Briefly, worms were transferred to assay plates either directly from cultivation plates (‘naive’) or after 1 h of pre-exposure to 2-nonanone (‘pre-exposed’). Odour-avoidance behaviour was recorded for 12 min. Assays were conducted on 90 mm agar plates, with the plate centre defined as (*x, y*) = (0 mm, 0 mm). Two odour sources (30% 2-nonanone in ethanol) were placed at (−22.5 mm, +15.0 mm) and (−22.5 mm, −15.0 mm), and worms were initially placed near the centre. Worm movement was recorded at 2 frames per second (fps)[14], and videos were processed using Move-tr/2D (v8.31; Library Inc., Japan) to extract centroid (*x, y*) coordinates. For the analyses presented here, we focused on the 2–7 min window of the assay, during which avoidance behaviour changes rapidly over time and space. The first 2 min were excluded because avoidance is typically not robust during this period, likely owing to the rapid increase in odour concentration[5, 41]. In total, 162 trajectories from pre-exposed worms and 163 trajectories from naive worms were obtained (the data split for deep learning is described below; 33 naive and 33 pre-exposed trajectories served as the test set and were also used for the post hoc analysis). Trajectories were down-sampled to 1 fps, yielding 300 frames (5 min × 60 s × 1 fps) of centroid *x*–*y* coordinates per trajectory.

#### 4.1.2 Courtship behaviour in fruit flies

A wild-type strain of *D. melanogaster* was used for the experiments. Ethical approval was not required for this data collection. Adult males and females were collected within 6 h of eclosion. To obtain virgin flies, males were maintained individually and females in groups of approximately 20–40 individuals. To obtain mated females, females were maintained in a group of approximately 20 males and 20 females. Flies were kept at 25 ^◦^C in 40–60% relative humidity under a 12 h light/dark cycle. Flies aged 6–10 days after eclosion were used in the following experiments.

To observe courtship behaviour, a male and a female were introduced into a chamber with sloped walls (60 mm in diameter, 3.50 mm in depth; modified from [42]). Videos were recorded with a CMOS camera (DFK 33UP1300, The Imaging Source Asia Co., Ltd.) equipped with a zoom lens (M0814-MP2, CBC Optics Co., Ltd.). Recording was initiated immediately after transfer of the flies and continued for 30 min at 30 fps. Videos were processed using Caltech FlyTracker to obtain fly centroid *x*– *y* coordinates and maximum apparent wing angles (‘max wing ang’)[19]. In total, 11 videos were collected for the virgin-female–virgin-male condition and 8 for the mated-female–virgin-male condition. To enable deep-learning-based exploratory analysis with a limited number of videos, each recording was segmented into 90 s clips; clips recorded after the onset of copulation were excluded to focus on courtship behaviour. This yielded 180 clips for the virgin-female–virgin-male condition and 149 for the mated-female–virgin-male condition (the data split for deep learning is described below; 36 virgin-female–virgin-male and 31 mated-female–virgin-male clips served as the test set and were also used for the post hoc analysis). Each clip comprised 2,700 frames (90 s × 30 fps) of centroid *x*–*y* coordinates and maximum apparent wing angles for two flies.

#### 4.1.3 Sex-dependent predator-avoidance strategies in scallops

The experimental subjects were Japanese scallops (*Mizuhopecten yessoensis*) and their predator, sea stars (*Asterias amurensis*). Ethical approval was not required for this data collection under current research guidelines. Nevertheless, all animals were handled with care to minimise stress and were maintained under appropriate conditions throughout the experimental period. Animals were collected using dredge nets in November 2023 and April 2024 from scallop fishing grounds off Monbetsu and Abashiri in eastern Hokkaido, Japan. All experiments used non-reproductive adult scallops (shell length *>* 70 mm) and sea stars capable of preying on scallops (arm-tip-to-arm-tip diameter *>* 130 mm). After collection, the animals were transported to the laboratory facilities of the Hokkaido Research Organization, Central Fisheries Research Institute, and acclimatised before experiments began.

Each scallop was individually marked on the left valve using underwater adhesive and stickers to enable identification throughout the experiments. Sex was determined by briefly inserting an endoscopic camera into the shell cavity and observing gonad colouration (creamy white in males and reddish-pink in females). To test how sex composition affected scallop behaviour in the presence of a predator, three experimental conditions were established: a male condition (ten male scallops with one sea star), a female condition (ten female scallops with one sea star), and a mixed condition (five male and five female scallops with one sea star). Experiments were conducted during the non-reproductive season (November–February) to avoid behavioural effects associated with reproduction. Ten trials were performed for each treatment, and each trial lasted 72–120 h.

Experiments were conducted in a circular tank 194 cm in diameter (radius 97 cm), with a bottom area of approximately 3 m^2^. Filtered seawater was supplied at 190 L h^−1^ and maintained at 10–12 ^◦^C under controlled lighting (300–500 lx). No food was provided during the experiments. Animal densities were set to three scallops per m^2^ and 0.3 sea stars per m^2^, corresponding to moderate densities observed in natural habitats[21]. All procedures followed the methodology established by Miyoshi[21] for scallop–sea star predation experiments.

Behaviour was recorded using a video camera mounted vertically above the tank. The camera was positioned to capture the entire tank bottom in a single frame and was programmed to record time-lapse images at 5 s intervals (12 frames per minute) throughout the 72–120 h period. Videos were analysed with DeepLabCut[1] to track the centre point of each individual and generate time-series *x*–*y* coordinate data. To enable deep-learning-based analyses with a relatively small number of videos, each 72–120 h video was segmented into 5 h clips, and the first 5 h were excluded as an acclimatisation period. Tracking errors (displacements *>* 99.96 cm between consecutive frames) were corrected by replacing the coordinate with the previous value. Clips with more than 100 such corrections were considered excessively noisy and were excluded. This procedure yielded 115 trajectory clips from the male condition, 143 from the female condition, and 157 from the mixed condition (the data split for deep learning is described below; 23 male, 30 female, and 32 mixed clips served as the test set and were also used for the post hoc analysis). Each clip contained 3,600 frames (300 min × 12 frames per min) of *x*–*y* coordinates for the body centre.

Several limitations of the experimental set-up should be acknowledged. First, in the confined circular tank, it was sometimes difficult to distinguish deliberate predatory searching by sea stars from random encounters with scallops. Second, the smooth tank bottom may have limited scallop movement, particularly during escape responses, precluding analysis of the scallops’ escape direction. Third, scallops occasionally attached to the tank wall, placing them slightly beyond the nominal tank radius in the coordinate system. Finally, even when animals appeared largely stationary, the tracking system detected small movements (approximately 0.33 cm min^−1^, equivalent to 100 cm over a 5 h clip). These movements likely reflect a combination of fine postural adjustments and system-level noise caused by water-surface motion, but they affected all conditions equally.

#### 4.1.4 Activity in isolated ants

The original dataset was collected by Koto[28] using the ant *Camponotus fellah*. Ethical approval was not required for this data collection.

Minor worker ants (body size *<* 8 mm; age *>* 7 months) were separated from their colony and housed for 24 h in a plastic box (105 mm × 87 mm) containing food and water (10 mm in diameter) and a light-shielded nest box (28 mm in diameter). Melatonin was added to the drinking water at a concentration of 1 µg mL^−1^. Ants were tagged with unique matrix codes (1.6 mm × 1.6 mm) and tracked at 2 fps using software developed by Mersch[43]. Data from the first 30 min of the experiment were discarded for all ants. Missing values were handled as follows. Gaps lasting ≤ 4 consecutive frames were linearly interpolated. Gaps lasting ≥ 5 consecutive frames were interpolated only when the distance between the last detected position before the gap and the first detected position after the gap was ≤ 3.75 mm; otherwise they were retained as missing values (NaN). A ‘trip’ was defined as the trajectory from the moment an ant left the nest until it re-entered the nest. For each trip, segments separated by missing data were treated as ‘sub-trips’. All coordinates were recorded in pixels and converted to metric units using a scaling factor of 0.05 mm per pixel, based on the arena dimensions (105 mm × 87 mm corresponding to 2,112 × 1,722 pixels). Ants that died during the experiment were excluded from the following procedure (6 isolated ants and 2 melatonin-treated ants).

After processing data from 27 isolated ants and 32 melatonin-treated ants (excluding dead individuals), the total number of sub-trips was 22,148 (isolated; control) and 24,708 (melatonin-treated). Of these, 19,269 (control) and 21,626 (melatonin-treated) had durations ≤ 60 s. This predominance of short sub-trips likely reflects a limitation of matrix-code tagging: tracking can be interrupted when an ant changes posture. Conversely, extremely long sub-trips lasting more than 300 s (control: 657 sub-trips; melatonin-treated: 556) may include trajectories in which ants remained stationary in the same location for extended periods. Accordingly, sub-trips with time-series lengths of 60–300 s were selected for analysis (control: 2,222 sub-trips; melatonin-treated: 2,526; the data split for deep learning is described below; 445 control and 506 melatonin-treated sub-trips served as the test set and were also used for the post hoc analysis). Each sub-trip consisted of 120–600 frames (60–300 s × 2 fps) of *x*–*y* coordinates.

#### 4.1.5 Behaviour of a rat model of schizophrenia

The original dataset was collected by Mundorf[30] in an open-field test using adolescent Sprague–Dawley rats in a schizophrenia model. All procedures complied with the German Animal Welfare Act and were approved by LANUV (Landesamt für Natur, Umwelt und Verbraucherschutz North Rhine-Westphalia).

Maternal immune activation (MIA) induced with Poly I:C is a well-established method for producing schizophrenia-like phenotypes[44]. Pregnant rats received 10 mg kg^−1^ Poly I:C (polyinosinic:polycytidylic acid dissolved in 1 mL of 0.9% NaCl) on gestational day 15, while control rats received 1 mL kg^−1^ of 0.9% NaCl. On postnatal day 39, 13 control offspring (5 females, 8 males) and 18 MIA offspring (7 females, 11 males) were tested in a 100 cm × 100 cm open-field arena for 5 min. Videos were recorded at 25 fps and processed with DeepLabCut[1] to extract *x*–*y* coordinates for 11 body parts: left ear, right ear, head, left shoulder, right shoulder, middle shoulder, body centre, left leg, right leg, middle leg, and tail. Tracking errors (body-part displacements *>* 31.25 cm between consecutive frames) were corrected by replacing the affected coordinate with the previous value. Four MIA rats (2 males, 2 females) were excluded owing to recording issues (videos recorded at *<* 25 fps), giving a final sample of 13 control rats (5 females, 8 males) and 14 MIA rats (5 females, 9 males).

To enable deep-learning-based analyses with a relatively small number of videos, each 5 min recording was segmented into 25 s clips. The first and last 12.5 s of each video were discarded, leaving 275 s for analysis (11 clips per rat). This yielded 143 trajectory clips from control rats and 154 from MIA rats (the data split for deep learning is described below; 30 control and 32 MIA clips served as the test set and were also used for the post hoc analysis). Each clip comprised 625 frames (25 s × 25 fps) of *x*–*y* coordinates for all 11 body parts.

#### 4.1.6 Social relationship formation in rats

All procedures were conducted in accordance with the guidelines of the University of Tokyo and were approved by the Animal Experiment Committee of the University of Tokyo, Graduate School of Arts and Sciences (Approval No. 2022-5).

Subjects were 32 male Sprague–Dawley (SD) rats, purchased from SLC Japan at 8 weeks of age. Animals were initially pair-housed in randomly assigned pairs. At the onset of group housing (approximately 12 weeks of age), eight groups of four rats were formed by combining two pairs. Before group housing, the animals underwent a battery of behavioural tests (glove test, open-field test, novel-object test, elevated-plus-maze test, and social-ranking test) to assess behavioural traits. These test data were beyond the scope of the present exploratory analysis and are not analysed here.

Group housing was conducted in a large custom-built cage consisting of a black aluminium frame, a black acrylic floor, and black polypropylene walls (85 cm × 85 cm × 155 cm [height]; inner dimensions 80 cm × 80 cm). A water-access opening was located 15 cm above the floor at the centre of the front wall. The floor was covered with recycled paper bedding. For identification, each rat was marked with one of four colours (blue, red, black, or orange) using animal-safe, non-toxic dyes. The marking pattern consisted of a belt-like line around the body and a line extending from the upper back to the base of the tail. During group housing, subjects remained in the same cage for three weeks. Daily tracking sessions were conducted for 12 h (19:00–07:00), corresponding to the entire dark phase. Dim indirect lighting (approximately 11 lx at the centre of the cage floor) was provided, and behaviour was recorded at 24 fps. Water and food (scattered around the centre of the cage) were provided daily. To maintain stable identification, markings were reapplied every three days; on the same days, bedding was partially replaced. Videos were processed using DeepLabCut[1], and *x*– *y* coordinates were extracted for eight body parts (nose, forehead, left ear, right ear, neck, upper back, lower back, and base of the tail) in each frame.

Data from days 2–3 of week 1 and days 20–21 of week 3 were selected for analysis. These days followed re-marking and cleaning days (days 1 and 19) to minimise potential effects of experimenter intervention. Each 12 h recording was segmented into 5 min clips to enable deep-learning-based analyses with a limited number of videos. To focus on periods of dynamic social interaction, we retained clips in which all rats travelled at least 240 cm (equivalent to three cage lengths) within the clip. To avoid sampling bias, 70 trajectory clips were randomly sampled from each of the eight groups, yielding 560 clips for the week-1 condition and 560 for the week-3 condition (the data split for deep learning is described below; 112 week-1 and 112 week-3 clips served as the test set and were also used for the post hoc analysis). Each clip comprised 2,400 frames (300 s × 8 fps after downsampling) of upper-back *x*–*y* coordinates, because this point was tracked most reliably.

#### 4.1.7 Behaviour of a mouse model of depression

All experimental procedures were approved by the Doshisha University Animal Care and Use Committee (Approval No. A25068).

To generate behavioural data for DiverScan analysis, mice underwent open-field testing under two conditions: a phosphate-buffered saline (PBS) control condition and a pharmacologically induced depression-like condition. The depression-like condition was induced using lipopolysaccharide (LPS; L-4130, serotype O111; Sigma-Aldrich), a well-established pharmacological agent for inducing depression-like behavioural phenotypes in rodents[45]. Hereafter, the LPS-induced depression-like condition is referred to as the ‘depression model’, whereas the PBS condition is referred to as the ‘control’.

Twenty-seven genetically unmodified mice with a C57BL/6J genetic background, aged 10–18 months, were used, comprising 9 males and 18 females. Mice were housed in same-sex groups of three per cage, giving three male cages and six female cages. Each mouse was tested under both the control and depression-model conditions. For each test, mice received an intraperitoneal (i.p.) injection of either LPS (0.01 mg kg^−1^) dissolved in PBS or an equivalent volume of PBS alone. The LPS dose was selected to induce depression-like behavioural changes while avoiding complete immobility during behavioural testing.

Before the injection–test cycles, habituation trials were conducted without PBS or LPS administration for all possible pairwise combinations of home cages. Each cage pairing underwent one 1 h habituation session. Because the same mice were subsequently tested repeatedly, habituation was performed to familiarise them with both the open-field apparatus and the mice from the paired home cage. This procedure was intended to minimise the influence of novelty associated with the apparatus and cage pairing on behaviour, allowing differences observed between the subsequent control and depression-model conditions to be interpreted primarily in relation to the treatment condition.

Following habituation, data were collected in a fixed sequence. Two PBS injection– test cycles were completed before any exposure to LPS, followed by two LPS injection–test cycles. Each cycle began with the relevant injection, and open-field testing was conducted 24–36 h later. Thus, the two cycles under each condition followed separate administrations. PBS testing was completed before any exposure to LPS because complete recovery to the original physiological and behavioural state after LPS administration could not be guaranteed.

In each injection–test cycle, all possible pairwise combinations of home cages were tested. Each open-field session included the six same-sex mice housed in a given pair of cages (three mice per cage). The three male cages yielded three unique cage combinations, whereas the six female cages yielded 15 unique combinations. Male and female mice were tested in separate sessions, and each session consisted exclusively of mice under either the control or the depression-model condition. Thus, each cycle comprised 18 recording sessions: 3 male sessions and 15 female sessions. Overall, 36 sessions were recorded under each condition, comprising 6 male sessions and 30 female sessions.

The open-field apparatus was placed inside a soundproof enclosure and consisted of a circular white acrylic floor 40 cm in diameter surrounded by a transparent acrylic wall 25 cm in height. In each session, the six mice were placed together in the arena, and their behaviour was recorded for 30 min at 30 frames per second from directly above the centre of the arena using an overhead camera.

The recorded videos were processed using vmTracking[46, 47], a DeepLabCut-based tracking method[1, 4]. For each mouse, vmTracking extracted the *x*–*y* coordinates of eight body parts: the nose, left ear, right ear, head, neck, body centre, body rear, and tail base.

To enable deep-learning-based analyses with a relatively small number of videos, each 30 min recording was segmented into 5 min clips, and the first 5 min of each video were discarded. This yielded 180 trajectory clips from the depression (LPS) condition and 180 from the control (PBS) condition (the data split for deep learning is described below; 36 control and 36 depression clips served as the test set and were also used for the post hoc analysis). Each clip comprised 3,000 frames (300 s × 10 fps after downsampling) of *x*–*y* coordinates for all eight body parts.

### 4.2 Step 1: Dataset construction

#### 4.2.1 Movement and user-defined input features

DiverScan accepts CSV files provided in the DeepLabCut format and/or our own format (user-defined time-series data). A default set of basic movement features (marked with an asterisk below) is selected automatically; through the checkboxes of the graphical user interface (Extended Data Fig. A2b), the user can add further features or remove any of the defaults. The selected features are then computed automatically as a time series 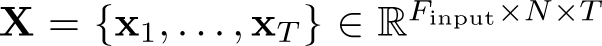, where 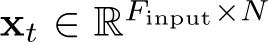. Here *F*_input_ is the number of movement features selected (e.g., speed, turning angle), *N* is the number of nodes, and *T* is the number of time steps. In the single mode, *N* = 1 always. Features marked with an asterisk (*) are selected by default.

### • *x* coordinate

- *y* **coordinate** In both the single and multi modes, the CSV file may include one or more tracked body parts per animal. DiverScan uses the *x*–*y* coordinates of a user-selected body part as the animal’s primary position, or it computes the animal’s centroid from the selected body parts and uses that as the primary position. This primary position is then used to calculate other features, such as speed and turning angle.
- **• Speed\***
- **Acceleration**
- **Azimuth\*** Azimuth is an absolute travel angle measured clockwise from the *y*-axis (north).
- **Turning angle\*** The turning angle is the change in travel direction relative to the previous time step.
- **Distance from the user-defined location** The distance from the user-defined location is calculated using user-provided *x*–*y* coordinates (e.g., the centre of the arena). In this study, we used (0, 0), the nest entrance, for the ant dataset, and the centre of the arena for the other datasets.
- **Angle from the user-defined location**
- **Distance from the initial location\***
- **Angle from the initial location\***
- **Cumulative travel distance**
- **Cumulative change in turning angle** The cumulative change in turning angle is the running total of the step-to-step changes in turning angle over time. If the turning angle stays the same, the cumulative change is zero.
- **Head direction** If an animal has two or more body parts, the user can identify which part is the head and which is the tail. Based on this, the azimuth of the animal’s head (i.e., its heading direction) is calculated.
- **User-defined additional features** The user can prepare additional features of interest (e.g., apparent wing angle, species).

#### 4.2.2 Pose represented by angles between body parts

In the single or multi mode with three or more body parts, DiverScan represents an animal’s pose using the angles defined by pairs of body parts and the centroid of all body parts: 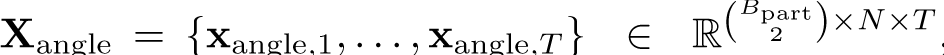, where *B*_part_ is the number of body parts and 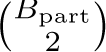 is the number of body-part pairs. For example, for an animal with three body parts A, B, and C, DiverScan computes, at each time step, the angles formed by A–centroid–B, A–centroid–C, and B–centroid–C. If the user has pose features generated by other software, these can be provided to DiverScan as ‘user-defined additional features’ (as described above).

#### 4.2.3 Adjacency matrix and edge features (multi mode)

In the multi mode, the adjacency matrix encodes relationships between nodes. This matrix is time-varying, and its weights change over time: **A** = {a_1_, . . ., a_*T*_} ϵ ℝ^*F*edge×*N*×*N*×*T*^.

In this study, we use a bidirectional complete graph to explore the data without imposing any a priori assumptions, although graphs with arbitrary structures are also supported. Let *V* = {1, 2, *. . ., N*} be the set of nodes. The edge set *E* for a bidirectional complete graph is defined as *E* = {(*i, j*) ∈ *V* × *V* | *i ≠ j*}. Thus, for every pair of distinct nodes (*i, j*), there is a directed edge from *i* to *j*, yielding a total of *N* (*N* − 1) directed edges.

Given a graph structure, DiverScan automatically computes the corresponding weighted adjacency matrix. Asterisks (*) mark the features selected by default. The user can add further features or remove any of the defaults:

- **Distance between nodes\***
- **Azimuth difference between nodes**

The difference in movement direction between two nodes. Smaller values indicate that the two animals are moving together.

### • Head-direction difference between nodes

The difference in head direction between two nodes. Smaller values indicate that the two animals are facing the same direction.

### • Body-part-to-body-part distance between nodes

In the multi mode, the user can specify pairs of body parts (e.g., body part A–body part B), and DiverScan computes the distance between the corresponding body parts of two animals (e.g., the distance between body part A of animal Y and body part B of animal Z).

#### 4.2.4 Features used for each dataset

For each dataset, the movement features used as input were selected from the lists above; the full selection for every dataset is given in Supplementary Table 1. In the single mode, these features were used directly, whereas in the multi mode they were organised as node and edge features of the time-series graph. Unless otherwise noted, the user-defined location was set to the centre of the arena; for the ant data it was set to the nest entrance (0, 0). Two datasets used an additional user-defined node feature: the apparent wing angle for the fruit-fly data and the species identity (scallop or sea star, encoded as a binary value) for the scallop–sea star data.

For the scallop–sea star data, we deliberately excluded features that directly encode absolute position in the tank, such as the distance and angle from the user-defined location (set to the tank centre; Supplementary Table 1). This let DiverScan capture scallop and sea star dynamics without relying on their absolute positions. Because scallops and sea stars spend much of their time stationary, absolute position information could leak across the training/validation/test splits of clips; however, the absolute location of any particular individual does not contribute to this study’s knowledge-discovery goal. Relative positions between individuals (e.g., inter-animal distance) could also leak across splits, but we consider this information meaningful for the present question and therefore retain it: our aim is not classification but exploratory data analysis focused on knowledge discovery.

#### 4.2.5 Label for classification

The user creates one folder for each experimental condition and stores the corresponding CSV files in it. Based on this folder structure, DiverScan automatically generates ground-truth labels for each condition using one-hot encoding. Hereafter, we refer to the experimental conditions as ‘classes’.

#### 4.2.6 Data splitting, standardisation, and zero-padding

DiverScan randomly splits the trajectories of each class into a training set (60%), a validation set (20%), and a test set (20%). Each feature fed into DiverScan-Net is standardised by z-score normalisation using the mean and standard deviation computed from the training set. If the prepared trajectories vary in time-series length, zero-padding is applied to match the length of the longest trajectory. The training and validation sets are used for learning, whereas the test set is held out and used independently for evaluation and post hoc analysis, thereby avoiding overfitting as far as possible.

We note an important limitation concerning the independence of the analytical units. In several datasets, multiple clips are obtained from the same individual; more generally, even samples that are not split into clips may originate from the same experiment, colony, or recording batch. The samples used for analysis should therefore not be regarded as fully independent. Consequently, even with the held-out design above, the samples used for training are not fully independent of those used for evaluation and post hoc analysis; that is, both stages are affected by pseudoreplication rather than operating on strictly independent samples. This is an inherent limitation of the Diver-Scan framework: because the dependence is introduced at the deep-learning stage, it cannot be resolved by the post hoc analysis alone (for example, by introducing a mixed-effects model). Despite this limitation, we consider it valuable to provide a new methodology that proposes hypotheses to researchers for the fine-grained time-series analysis of animal movement—a field in which behavioural researchers have long been constrained in practice (see Discussion).

### 4.3 DiverScan-Net for Steps 2–4

#### 4.3.1 Hardware and software for machine learning

We ran experiments on both a standard Google Colab GPU instance (Tesla T4, 15 GB GPU memory, 12.7 GB system RAM) and a local GPU server (NVIDIA RTX A6000; AMD EPYC 7313P CPU). For training, we used the versions of PyTorch and CUDA provided by the Google Colab environment.

#### 4.3.2 Architecture of DiverScan-Net

The input of DiverScan-Net is a time series 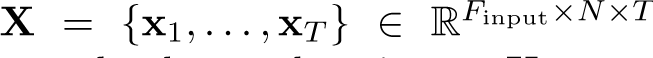. The time series of angles between body parts can also be used as input: **X**_angle_ = 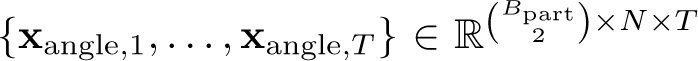, where *B*_part_ is the number of body parts. In the multi mode, a temporally dynamic weighted adjacency matrix is additionally used as input: 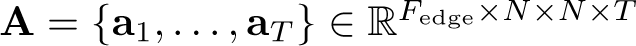.

The user specifies the number of attention branches *B* in DiverScan-Net, and the network splits into *B* branches accordingly. The output of DiverScan-Net is the predicted class for each attention branch: **Y** = {**y**_1_, *. . .,* **y***_B_*} ∈ R*^B^*^×|Class|^. An example network is shown in Extended Data Fig. A9a.

Note that DiverScan-Net preserves the time-step dimension *T* and node dimension *N* all the way through to the linear block. This design avoids compressing temporal and node information and thus maintains spatio-temporal resolution. Consequently, the attention time series (attention map) computed in the attention block retains the original temporal length and node dimension, which may enhance interpretability: **Attn** = {**attn**_1_, *. . .,* **attn***_T_*} ∈ R*^N^*^×^*^T^* for each attention branch.

Details of the individual blocks in DiverScan-Net are described below.

##### • Linear block for angles between body parts

The input to this block is a time series of angles between body parts, **X**_angle_ ∈ 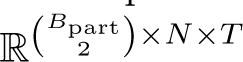. This block reduces the 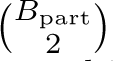 dimension to a single channel: **H***_angle_* ∈ R^1×^*^N^*^×^*^T^ .* When DiverScan-Net receives this matrix as one of its inputs, exactly one instance of this block is inserted immediately after the input (Extended Data Fig. A9b). Its output **H**_angle_, with shape (1, *N, T*), is then concatenated with the input **X**, with shape (*F*_input_, *N, T*).

##### • Feature-extraction block (single mode)

This block takes as input either 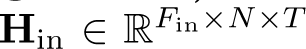 from the previous block, or the raw input 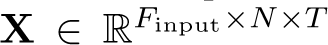 if it is the first block in DiverScan-Net. The block applies either a one-dimensional temporal convolution or a bidirectional LSTM; the choice is automatically determined by the neural architecture search (see below). The output is 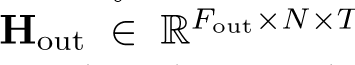. In the single mode, *N* = 1 always, so this dimension is omitted in the actual computation; we retain the *N* -dimensional form in the notation only for consistency with the multi mode. When the one-dimensional convolution is selected, it is applied along the *T* dimension, with a residual (skip) connection implemented within the block by summation. When the bidirectional LSTM is selected, it learns temporal dependencies in the time series, and a dropout layer is applied after it.

##### • Feature-extraction block (multi mode)

This block takes two inputs. The first is either 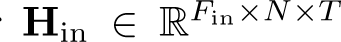 from the previous block, or the raw input 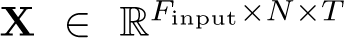 if it is the first block. The second is the temporally dynamic adjacency matrix 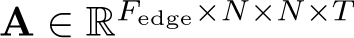. The block applies a convolution based on the spatio-temporal graph convolutional network (ST-GCN)[48], producing 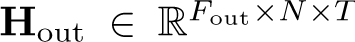. First, a one-dimensional convolution with temporal kernel size *t*_kernel_ and spatial size 1 is applied to **H**_in_ or **X** to extract temporal features. In addition, learnable edge weighting is introduced: a linear layer computes trainable weights for each edge, which are multiplied with the precomputed normalised adjacency matrix. The graph convolution is expressed as an Einstein summation that aggregates the weighted contributions of neighbouring nodes. Specifically, the output feature at batch index *n*, channel *c*, time step *t*, and node *w* is *y_nctw_* = ∑*_k,v_ x_nkctv_ A_nkvwt_,* where *k* denotes the spatial kernel dimension and *v* the input node index. After the graph convolution, the feature map is processed by batch normalisation, a ReLU activation, a further temporal convolution to refine the temporal dynamics, and a residual connection within the block.

##### • Attention block

This block takes as input either 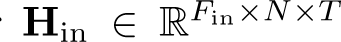 from the previous block, or the raw input 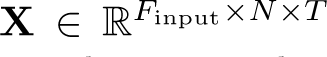 if it is the first block. It applies a one-dimensional convolution with temporal and spatial kernel sizes both equal to 1 to obtain an attention time series, **Attn** ∈ R*^N^*^×^*^T^ .* A softmax then normalises the attention values to lie in [0, 1] and sum to 1: over the time axis in the single mode, and jointly over all time points and nodes (a global softmax over the *N* × *T* entries) in the multi mode. The output of the block is the Hadamard (element-wise) product of the input and the attention time series, 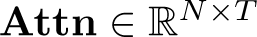, so that the subsequent block receives an attention-weighted feature matrix.

##### • Linear block

This block takes as input 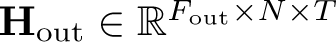 from the previous block and applies a linear layer to predict the class of the time series. In each branch, exactly one instance of this block is placed immediately before the output. Optionally, the user can shuffle the data along the *N* dimension during training only, prior to the linear layer, so that training does not rely on any specific node (i.e., any specific individual) in the multi mode. In this study we applied this option to all multi-mode datasets except the fruit-fly and schizophrenia-model rat datasets, because in these two the nodes play asymmetric, fixed roles—male vs. female in flies, and distinct body parts of a fixed skeleton in the rat—so that shuffling nodes would destroy the role information that distinguishes them. The output of this block (i.e., the output of an attention branch) is **y** ∈ R^|Class|^.

#### 4.3.3 DiverScan loss for DiverScan-Net

DiverScan-Net is trained by minimising DiverScan loss, defined in Eq. (1). We present a general form that includes four components; individual components can be enabled or disabled by the user depending on the dataset (see below).

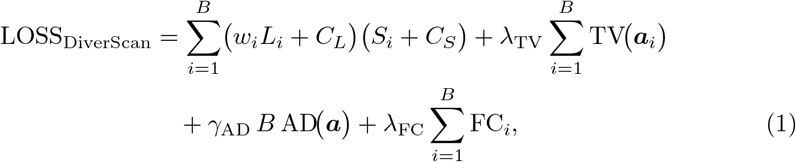

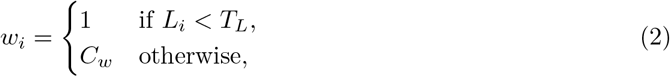

where *L_i_* is the cross-entropy loss of branch *i*, *S_i_*its attention-similarity score, ***a****_i_* its attention time series, and *B* the number of attention branches. The attention ***a****_i_* ∈ R*^N^*^×^*^T^* is defined over all *N* nodes and *T* time steps (*N* = 1 in the single mode). The equations below are written for the single mode; how each component treats the node axis in the multi mode is stated alongside it. The components are defined as follows.

The similarity *S_i_*rescales the cosine similarity between the attention time series of branch *i* and those of the other branches to the range [0, 1] and averages over them:

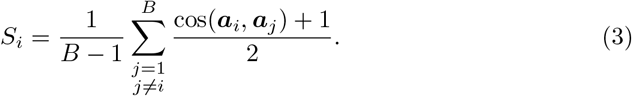

In the multi mode, the cosine similarity is computed over the flattened *N* ×*T* attention values, so that two branches count as dissimilar only when they attend to different node–time combinations, not merely to different nodes or different times alone.

The total-variation term measures the temporal smoothness of an attention time series:

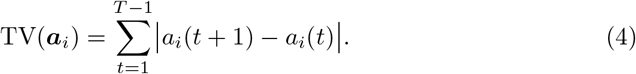

This difference is taken along the time axis; in the multi mode it is computed for each node separately and then summed over nodes, so that smoothness is encouraged in time but not across nodes.

The attention-distribution term penalises branches whose attention is excessively concentrated at both ends of the sequence:

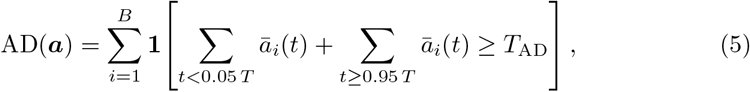

where *ā_i_*(*t*) is the batch-averaged attention of branch *i* and **1**[·] is the indicator function. The ends of the sequence are defined along the time axis; in the multi mode the attention in the first and last 5% of time steps is summed over all nodes. Because AD is built from an indicator function on the batch-averaged attention, it carries no gradient with respect to the attention weights; it therefore acts as a soft constraint that is not optimised directly during back-propagation but instead influences the neural architecture search (Step 2), which selects architectures with lower overall loss.

The feature-coherence term encourages the attended feature vectors to be consistent over time:

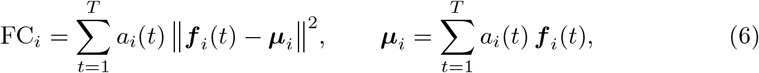

where ***f*** *_i_*(*t*) is the *D*-dimensional feature vector fed to the attention block of branch *i*. In the multi mode, the weighted mean ***µ****_i_* and the sum are computed for each node separately, so that the attended feature vectors are encouraged to be consistent over time within each node, without enforcing consistency across different nodes.

In Eq. (1), the first term sums, over branches, the product of the branch loss *L_i_*and the branch similarity *S_i_*. Because *L_i_* and *S_i_* are multiplied, *reducing both terms simultaneously yields a non-linear reduction in the loss*. To prevent the product from becoming excessively small, we add constants *C_L_*and *C_S_*(both ≥ 1) to *L_i_* and *S_i_*, respectively. The weight *w_i_* (Eq. 2) penalises a branch whose classification loss exceeds a threshold *T_L_*, with penalty magnitude *C_w_ >* 1; this discourages branches whose attention is dissimilar to the others but which do not classify well and therefore do not contribute to knowledge discovery. In the total-variation term, a smaller value corresponds to a smoother attention time series, discouraging the model from concentrating attention on only a few local time steps (e.g., isolated spikes) and thereby improving interpretability; *λ*_TV_ controls its strength. The attention-distribution term is scaled by *B* so that, as the number of branches grows, the constraint against end-concentrated attention becomes stronger, preventing many branches from collapsing their attention onto the very beginning or end of the sequence. The feature-coherence term encourages a branch to detect the same characteristic behaviour consistently whenever it occurs during the recording, which improves interpretability. It is optional and disabled by default (*λ*_FC_ = 0), because in animal behaviour the long-term context of when a behaviour occurs can itself be important—for example, when a characteristic behaviour arises specifically at the beginning or end of an experiment—and enforcing feature coherence could suppress such context-dependent structure. The user can enable it for datasets where this is not a concern.

##### Comparison with related approaches

A quantitative evaluation on synthetic signal data confirmed that each component of DiverScan loss contributes to training and performance, and that DiverScan loss outperforms related approaches (see Supplementary Note). Several prior studies have sought to diversify attention time series; for example, Li et al. added the negative cosine similarity between attention time series to the loss for language tasks[49, 50], so that lower similarity incurs a smaller penalty and each attention mechanism is encouraged to attend to different features. However, such approaches do not address three issues that arise for time-series data. First, different attentions can concentrate on nearby time steps, focusing on a very limited local subset that corresponds to the same event while still reducing the overall similarity. Second, attentions that are simply random also reduce the similarity. Neither case contributes to meaningful diversity. Third— and central to our goal of knowledge discovery—the interpretability of each attention is not directly related to model performance, such as the cross-entropy loss. DiverScan loss addresses these issues in a branching model, refining the attention time series in a way that is beneficial to knowledge discovery. Such meaningful diversity is enforced explicitly by DiverScan loss itself, rather than by the attention-branch architecture alone[51] (Supplementary Note).

##### Hyperparameter settings

For the animal datasets, we used *B* = 5, *C_L_*= 3, *C_S_*= 1, and *C_w_*= 10. (In the synthetic signal dataset, the classification and similarity constants are swapped, *C_L_* = 1 and *C_S_* = 3, placing relatively more weight on accurate classification; for the more complex animal data we instead placed more weight on attention separation. These constants are user-adjustable; see Supplementary Note.) For *T_L_* and the regularisation strengths, DiverScan sets default values based on the number of classes, |Class|:

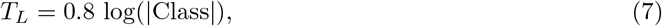

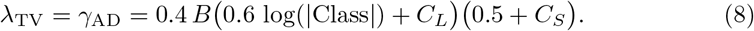

Equation (7) reflects that, for a balanced classification problem in which the model predicts at chance (probability 1*/*|Class| for every class), the cross-entropy loss equals log(|Class|); the factor 0.8 thus requires slightly better-than-chance performance. In Eq. (8), the term 0.6 log(|Class|) reflects the expected level of accurate classification and the term 0.5 the expected low similarity among attention time series, so that the first term of Eq. (1) maintains an approximately constant ratio to the regularisation terms as |Class| varies; the overall factor 0.4 balances the regularisation. By default, *γ*_AD_ takes the same value as *λ*_TV_. We used *T*_AD_ = 0.8.

### 4.4 Step 2: Neural architecture search

The timescale and structure of behaviours vary widely across animals and experimental set-ups, so a single fixed architecture is often not well suited to every dataset. DiverScan therefore optimises the network architecture automatically, searching for an architecture matched to the behaviours in each dataset, such as one with an appropriate temporal receptive field for each branch. Users who prefer not to run this search can instead use the predefined hyperparameters we provide. For the search we used Optuna[52] with the tree-structured Parzen estimator (TPE). For each animal dataset, 300–500 trials were run on the training and validation sets, and the best hyperparameters, which minimise the validation loss, are reported in Supplementary Table 1. To conserve computation, some hyperparameters were fixed across all trials rather than searched (e.g., the learning rate of the Adam optimiser, fixed at 0.001, and the batch size; see Supplementary Table 1). The linear block for angles between body parts and the linear block immediately before the output were not part of the search.

#### • Weight decay of the optimiser

The weight decay of the optimiser (Adam) is searched over 1 × 10^−5^ to 1 × 10^−1^ (log scale).

#### • Dropout probability

The dropout probability is searched over 0.1–0.5. To conserve computation, all dropout layers in DiverScan-Net share this probability.

#### • Feature-extraction blocks before branching

The number of blocks and the temporal kernel size are searched over 0–3 and 1– 10% of *T* (or of a user-defined length), respectively. When the number of blocks is 0, DiverScan-Net branches immediately after the input (Extended Data Fig. A9c). In this study, a user-defined length was set only for the ant dataset (121 frames, 20.2% of the maximum length), because the sub-trip length varies across samples in that dataset.

#### • Feature-extraction blocks in branches (single mode)

For each branch, we search over (i) the number of one-dimensional temporal convolution blocks (0–5), (ii) the temporal kernel size (1–10% of *T*, or of a user-defined length), and (iii) the number of LSTM blocks (0–5). Because each branch can have a different number of blocks and a different kernel size, the latent representations can differ across branches. Within a branch, the LSTM blocks are always placed before the temporal convolution blocks and the attention block.

#### • Feature-extraction blocks in branches (multi mode)

For each branch, the number of ST-GCN blocks and the temporal kernel size are searched over 0–7 and 1–10% of *T* (or of a user-defined length), respectively.

#### • Attention block

For each branch, exactly one attention block can be placed at any position (e.g., immediately after branching, between feature-extraction blocks, or immediately before the linear block), so that each branch can attend at the position best suited to the features it extracts. If a branch contains no feature-extraction blocks, it consists only of an attention block and a linear block.

Alternatively, the user can employ predefined hyperparameters that we have empirically found to perform well.

### 4.5 Step 3: Training

DiverScan-Net is trained using the hyperparameters obtained from the neural architecture search. For the animal datasets used in this study, the number of epochs was set to 100. Training used the training and validation sets, and the model weights from the epoch with the lowest validation loss were used in Step 4.

### 4.6 Step 4: Testing

The trained DiverScan-Net classifies the clips in the test set. For each clip, we save the attention map and the predicted class label from every branch. We report the classification performance of each branch as the macro-averaged accuracy:

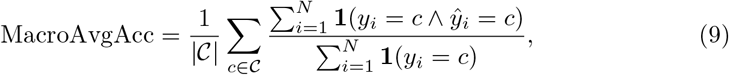

that is, the mean of the per-class accuracies, where C is the set of classes and *N* is the number of clips in the test set.

### 4.7 Step 5: Interpretation of attention

After DiverScan-Net has been tested (Step 4), the attention maps of the test clips are fed into the attention-interpretation module. This module supports the user— typically a specialist in the target animal—through two complementary submodules: visual inspection of the attention maps, and an LLM-based assistant that verbalises the captured differences. Throughout, a time bin is regarded as *highlighted* when its attention value exceeds the chance level 1*/*(*N T* ^′^), where *N* is the number of nodes and *T* ^′^ is the number of valid (non-padded) time steps.

#### 4.7.1 Visual inspection

For each animal’s trajectory, the segments to which DiverScan-Net attends are highlighted, giving the user an intuitive view of the model’s focus (Extended Data Fig. A2e). To make this quantitative, the module computes the correlation between each attention map and the predefined hand-crafted features (e.g., speed) and automatically ranks the features by how strongly they correlate with the attention, presenting the most positively and most negatively correlated features rather than requiring the user to inspect all of them (Extended Data Fig. A2e,h). An interactive plot of the trajectory is additionally provided for closer inspection (Extended Data Fig. A2g).

The module further visualises the diversity among attention branches using principal component analysis (PCA) of their attention maps (Extended Data Fig. A2f). This view removes the need to fine-tune the number of branches by hand: if the number of branches is set larger than the number of behavioural differences present in the data, branches that capture similar features simply appear close together in the PCA space, whereas branches capturing distinct differences are well separated. The user can therefore choose a generous number of branches and read off the effective diversity from the plot.

#### 4.7.2 LLM-based assistant

The second submodule uses an LLM to interpret and explain the behavioural difference captured by each attention branch, and to suggest its likely biological meaning. The LLM (gemini-3.1-pro-preview; run with temperature 0 and a fixed seed for reproducibility) generates its explanations from three sources of information.

The first is the correlation between the attention values and the predefined hand-crafted features, which addresses the question ‘when are the segments highlighted?’ (Extended Data Fig. A2h). These hand-crafted features comprise all the movement features listed in Step 1, regardless of whether they were selected as model input, together with their moving average and moving variance. In the multi mode, further quantities derived from the relations among nodes are added:

- the node difference of each movement feature (the sum of absolute differences of that feature between a node and all other nodes);
- from the *x*–*y* coordinates, the distance to the group centroid, the local density, and the head-direction alignment;
- edge features (e.g., the distance between animals) and their per-edge average and variance;
- the sum, minimum, maximum, and variance of the features of the edges connecting to each node;
- minimum-spanning-tree quantities (node degree, leaf indicator, and the sum and maximum of the tree edges connecting to each node);
- the number of clusters (connected components of the proximity graph obtained by thresholding the inter-node distance);
- and the spatial-configuration pattern (the proximity graph classified up to node relabelling, defined for up to five nodes).

The second is the difference in hand-crafted feature values between conditions within the highlighted segments—quantified by the correlation and the Hellinger distance—which addresses the question ‘what differs in the highlighted segments?’.

The third is a set of research papers relevant to the experiment: through retrieval-augmented generation (RAG), the LLM grounds its interpretation in literature that the user, as a specialist in the organism, considers trustworthy, thereby tailoring the explanation to the specific study and reducing ungrounded description. RAG is optional and can be disabled by the user.

Because the attention maps, their correlations with hand-crafted features, and the between-condition differences are all supplied to the LLM as a compact table, and the relevant literature is made available through RAG, the prompt itself can remain simple: the RAG-grounded LLM reasons largely on its own from this rich input. The initial prompt is generated automatically from a template, into which the user inserts only a brief description of the experiment. For example, the default prompt generated for the fruit-fly analysis was:

This file shows, for the courtship behavior of a pair of fruit flies, which hand-crafted features had high or low correlation with the attention (from an XAI model). Compare the characteristics of the [number of attention branches] attention branches and give me an overview. Also, explain the differences in experimental conditions across each of the attention branches.

CSV: [table of per-branch correlations and between-condition differences]

Because the module is realised as an interactive chat, the user can go beyond RAG and give the LLM explicit, case-specific instructions. This is useful because some of the automatically computed hand-crafted features may lack a clear meaning under a given analysis setting. In particular, features that increase monotonically over the full recording (such as accumulated travelled distance or elapsed time) can be meaningless once the recording is divided into clips, since each clip captures only a fragment of the original sequence; such features may nonetheless correlate with attention for reasons unrelated to the behaviour of interest. Similarly, attention that finds no discriminative signal in a given condition may default to the boundaries of the sequence, producing a spurious correlation with features that happen to rise at the edges. In such cases the user can instruct the LLM to down-weight or set aside the affected indicators, so that the interpretation prioritises features with genuine biological relevance; the visual-inspection and LLM submodules together make such artefacts easy to recognise and discount rather than take at face value. The chat also lets the user ask follow-up questions and request candidate post hoc analyses. In this way, human experts remain in control of context and biological relevance while DiverScan surfaces candidate differences for them to examine. Together with its support for time-series graphs, this attention-interpretation module allows DiverScan to provide rich, expert-guided insight into behaviour.

### 4.8 Post hoc analysis for scallop–sea star dataset

For the post hoc analyses, scallops that were preyed upon during the experiments (*N* = 15) were excluded to avoid behavioural biases (5 males in the male condition, 5 females in the female condition, and 4 males and 1 female in the mixed condition). For attention branch 5, we analysed non-overlapping 240 s segments that began when at least one scallop reached a speed of ≥ 15 cm min^−1^ (approximately twice the shell length per minute). Each segment was labelled by sea-star activity: if the sea star exceeded 15 cm min^−1^ at any point during the first 60 s of the segment, the segment was labelled ‘sea-star movement’; otherwise it was labelled ‘no sea-star movement’. For each segment we computed the mean displacement per scallop. Clips containing no segment with a given label were excluded from the corresponding comparison, so fewer clips entered the ‘sea-star movement’ comparison (male, 8; female, 6; mixed, 8) than the ‘no sea-star movement’ comparison (male, 22; female, 30; mixed, 32).

**Supplementary information. Supplementary Table 1: Input features and hyperparameters of DiverScan-Net for each dataset.**

Movement features used as input (directly in the single mode, or as node and edge features in the multi mode), and the hyperparameters explored by the neural architecture search (with the fixed ones also indicated), for every dataset. **Supplementary Table 2: Table supplied to the LLM-based assistant for the worm dataset.** Summary table supplied to the LLM-based assistant for interpretation (see Methods). Because an LLM’s ability to parse very large tables is limited, a practically abbreviated version was used (e.g., the top several and bottom several features for each measure). **Supplementary Table 3: Per-branch classification performance of DiverScan-Net.** Precision, recall, F1-score, and macro-averaged accuracy on the held-out test set for each dataset and each attention branch. **Supplementary Table 4: Details of the statistical tests.** Test used, sample sizes, test statistic, (adjusted) *p* value, 95% confidence interval, and effect size (where applicable) for every statistical comparison reported in the main text, figures, and Notes. **Supplementary Table 5: Table supplied to the LLM-based assistant for the fruit-fly dataset.** Provided for interpretation, with contents as described for Supplementary Table 2. **Supplementary Table 6: Table supplied to the LLM-based assistant for the scallop–sea star dataset.** Provided for interpretation, with contents as described for Supplementary Table 2. **Supplementary Table 7: Table supplied to the LLM-based assistant for the ant dataset.** Provided for interpretation, with contents as described for Supplementary Table 2. **Supplementary Table 8: Table supplied to the LLM-based assistant for the schizophrenia-model rat dataset.** Provided for interpretation, with contents as described for Supplementary Table 2. **Supplementary Table 9: Table supplied to the LLM-based assistant for the rat social-formation dataset.** Provided for interpretation, with contents as described for Supplementary Table 2. **Supplementary Table 10: Table supplied to the LLM-based assistant for the depression-model mouse dataset.** Provided for interpretation, with contents as described for Supplementary Table 2.

## Acknowledgements

M.Su. is supported by JSPS KAKENHI Grant Number JP23KJ1414, the Daiichi-Sankyo ‘Habataku’ Support Program for the Next Generation of Researchers (2nd round, No. 1), and the ‘Habataku’ Support Program for the Next Generation of Researchers at The University of Osaka (3rd round, No. 5). T.M. is supported by JSPS (KAKENHI JP25H01116 and JP21H05299) and JST CREST (JPMJCR23P2). K.D.K. is supported by JSPS (KAKENHI JP22KK0100 and JP25K02314), JST CREST (JPMJCR23B2), Grant-in-Aid for Research in Nagoya City University (2121101 and 2530002), and Nakatani Foundation. T.B. is supported by JST SPRING Grant Number JPMJSP2130. R.T. is supported by JSPS (KAKENHI JP22H05650 and JP24H01433). K.M. and Y.M. are supported by JSPS KAKENHI Grant Numbers JP21H05296, JP22H05663, JP24H01455. A.K. is supported by JSPS (KAKENHI JP21H05297) and AMED under Grant Number JP18gm6110014, and Suntory Rising Stars Encouragement Program in Life Sciences. M.Si. and H.N. are supported by JSPS KAKENHI Grant Number JP21H05297 and the MEXT Promotion of Distinctive Joint Usage/Research Center Support Program Grant Number JPMXP0724020292. H.M. is supported by JSPS KAKENHI Grant Number JP24K16867. S.H. is supported by JSPS KAKENHI Grant Numbers JP22KJ0702, JP24KJ0179. G.T. is supported by JSPS KAKENHI Grant Numbers JP24H01428, JP21H05175. K.O., N.K., and H.K. are supported by JSPS KAKENHI Grant Number JP23H05428. H.K. is supported by JSPS KAKENHI Grant Numbers JP21H04421, JP22H03914. H.A. is supported by JSPS KAKENHI Grant Number JP24K15711. S.T. is supported by JSPS (KAKENHI JP21H05296 and JP23H00502) and JST CREST (JPMJCR23P2). The illustrations of *C. elegans*, the fruit fly, the schizophrenia-model rat, and the depression-model mouse were created by Ms. Lamisa Mashiat, Ms. Haruka Ando, Prof. Yasunobu Yasoshima, and Dr. Kaoru Ide, respectively. The DiverScan logo was designed by M.Su., with invaluable help from Ms. Kana Yasuda. During the preparation of this manuscript, the authors used Claude (Anthropic) and ChatGPT (OpenAI) to assist with writing and programming. All such output was subsequently reviewed and edited critically by the authors, who take full responsibility for the content of the published article. The authors would like to thank all members of the authors’ laboratories for their valuable comments and support.

## Declarations

### • Ethics approval and consent to participate

Declared in Methods.

### • Consent for publication

Not applicable.

### • Data availability

The *C. elegans*, fruit-fly, ant, schizophrenia-model rat, and depression-model mouse datasets are available in our Zenodo repository (DOI: 10.5281/zenodo.22940681). The scallop–sea star and rat social-formation datasets are not publicly available but are available from the corresponding authors upon reasonable request.

### • Materials availability

Not applicable.

### • Code availability

Jupyter notebooks, source-code files, and user support are available in our GitHub repository (https://github.com/M-SUKEGAWA/DiverScan). A ready-to-use copy of DiverScan and demo notebooks are provided in a Google Drive folder (https://drive.google.com/drive/folders/1aW7cOEd50ctArG0MOgv-AJdWZFU-eDy5); instructions for their use are given in the GitHub repository. Full licence details can be viewed in the same repository.

### • Author contributions

M.Su. and T.M. conceived the project and designed Diver-Scan. M.Su. implemented DiverScan and analysed all data. For the *C. elegans* data, the experiment was designed by K.D.K. and S.J.Y., and data were collected by S.J.Y. For the fruit-fly data, the experiment was designed by R.T. and M.Su., and data were collected by R.T. For the scallop–sea star data, the experiment was designed by K.M. and C.Y.; the conceptual framework and analytical support were provided by Y.M.; sample collection was supported by C.Y.; and data were collected by K.M. For the ant data, the experiments were designed by A.K., and data were collected by A.K. Code for preprocessing behavioural data was developed by M.Si., T.I. and H.N. For the schizophrenia-model rat data, the experiment was designed by N.F. and A.M., and data were collected by A.M. For the rat social-formation data, the experiment was designed by S.H., N.K. and G.T., and data were collected by S.H. For the depression-model mouse data, the experiment was designed by H.A. and M.Su., and data were collected by H.A. The synthetic data were generated by M.Su. and T.M. The writing and discussion were carried out by all authors. H.A., T.B., N.F., S.H., T.I., N.K., K.D.K., H.K., A.K., Y.M., H.M., K.M., A.M., H.N., K.O., M.Si., S.T., R.T., G.T., C.Y. and S.J.Y. contributed equally to this work and are listed alphabetically in the author list.

## Appendix A Extended data figures

**Fig. A1.**
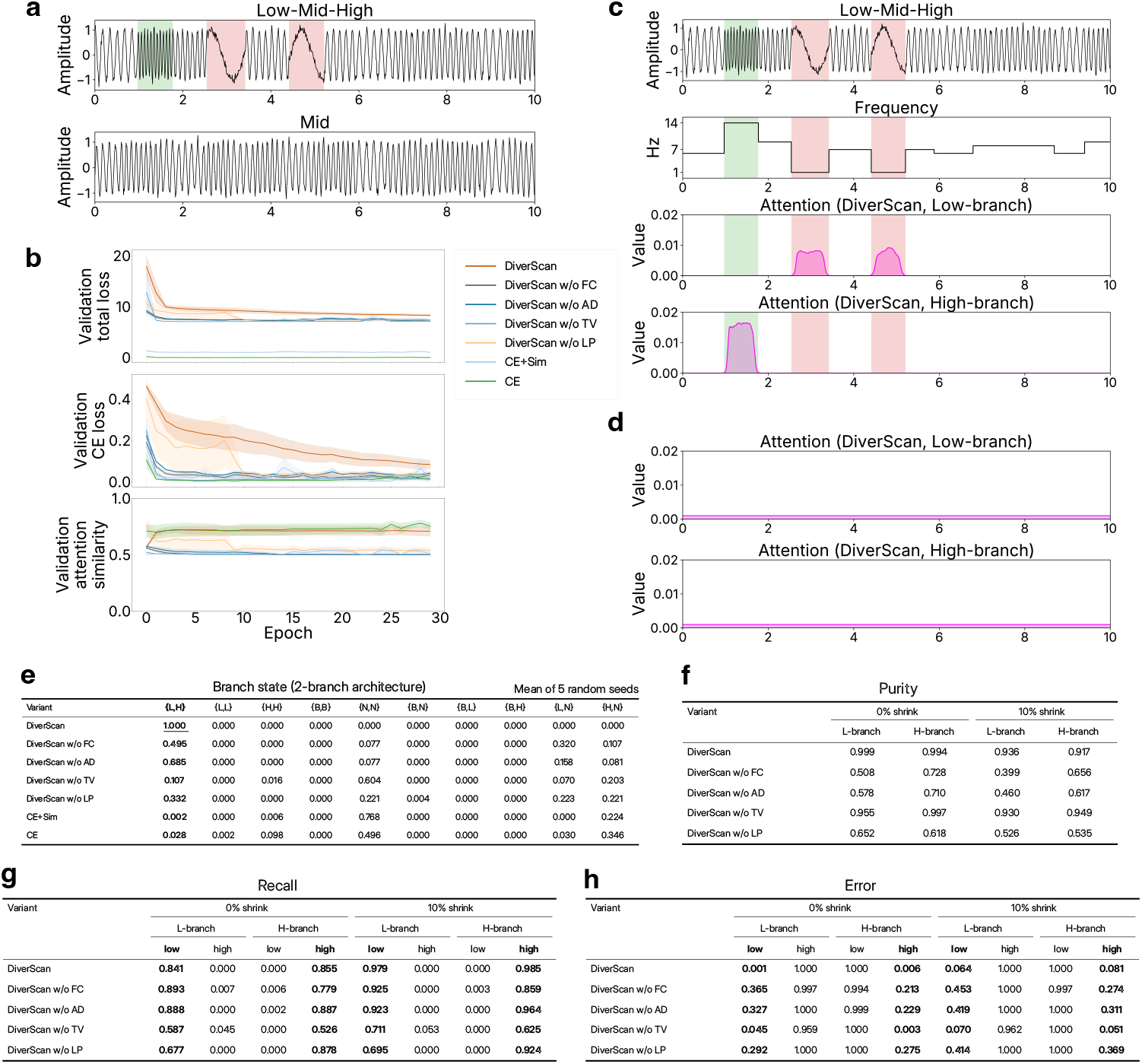
Quantitative evaluation of DiverScan loss using synthetic signals. See Supplementary Note for detailed discussion. **a,** Representative examples of the two signal conditions. Top: a Low-Mid-High signal containing low-, mid-, and high-frequency segments (the red and green shadings mark the low- and high-frequency segments, respectively). Bottom: a Mid signal containing only mid-frequency segments. **b,** Loss curves during training. From top to bottom: the total loss LOSS, the mean branch cross-entropy loss *L̄*, and the mean branch similarity *S̄* on the validation set, each shown as the mean SEM across the 5 random seeds. **c,** Attention time series produced by the DiverScan model for the Low-Mid-High signal shown in **a**. From top to bottom: the raw signal, the underlying frequency-of-segment time series, and the attention time series of the low-branch and the high-branch. **d,** Attention time series produced by the DiverScan model for the Mid signal shown in **a**. **e,** Distribution of per-sample branch states across correctly classified Low-Mid-High test samples. **f,** Purity of the highlighted time bins, computed on the subset of samples that reached the ideal *L, H* branch state. Values are the mean across the 5 random seeds (or fewer, when some seeds yielded no samples). **g,** Recall rate of the highlighted time bins. **h,** Error rate of the highlighted time bins.

**Fig. A2.**
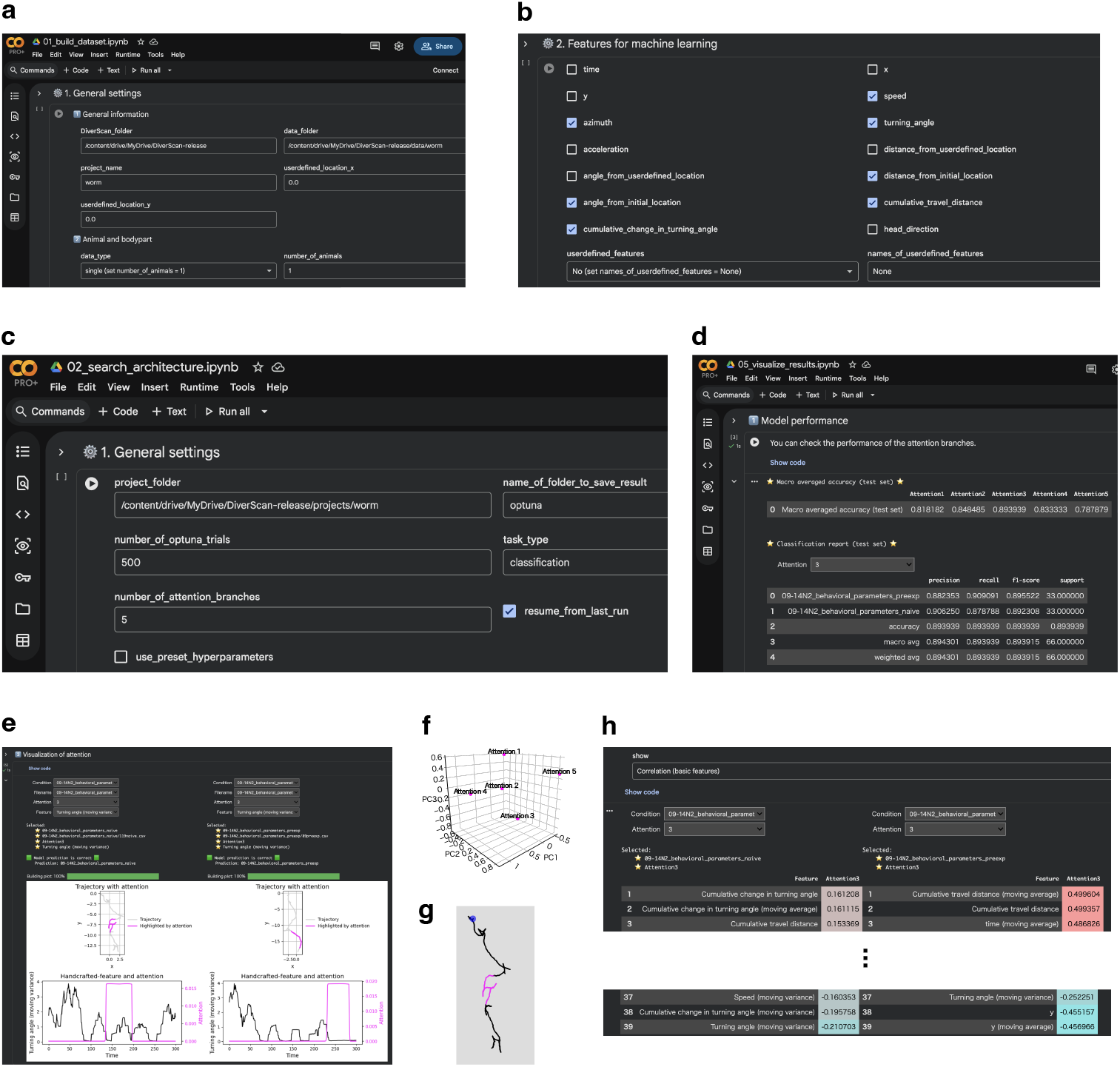
Graphical user interface (GUI) of DiverScan in a Jupyter notebook on Google Colab. **a**, Step 1: Simple settings for dataset construction. **b,** Step 1: Checkboxes for basic behavioural features used as input to DiverScan-Net. **c,** Step 2: Simple settings for architecture search. Steps 3 and 4 use similar GUIs. **d,** Step 4: Model performance. The results can be easily inspected in Step 5. **e,** Step 5: Attention maps visualised on trajectories and hand-crafted features. **f,** Step 5: PCA of the attention branches. **g,** Step 5: Interactive plot of a trajectory. **h,** Step 5: Hand-crafted features ranked by their correlation with the attention maps, so that the most strongly correlated features can be compared at a glance to aid interpretation.

**Fig. A3.**
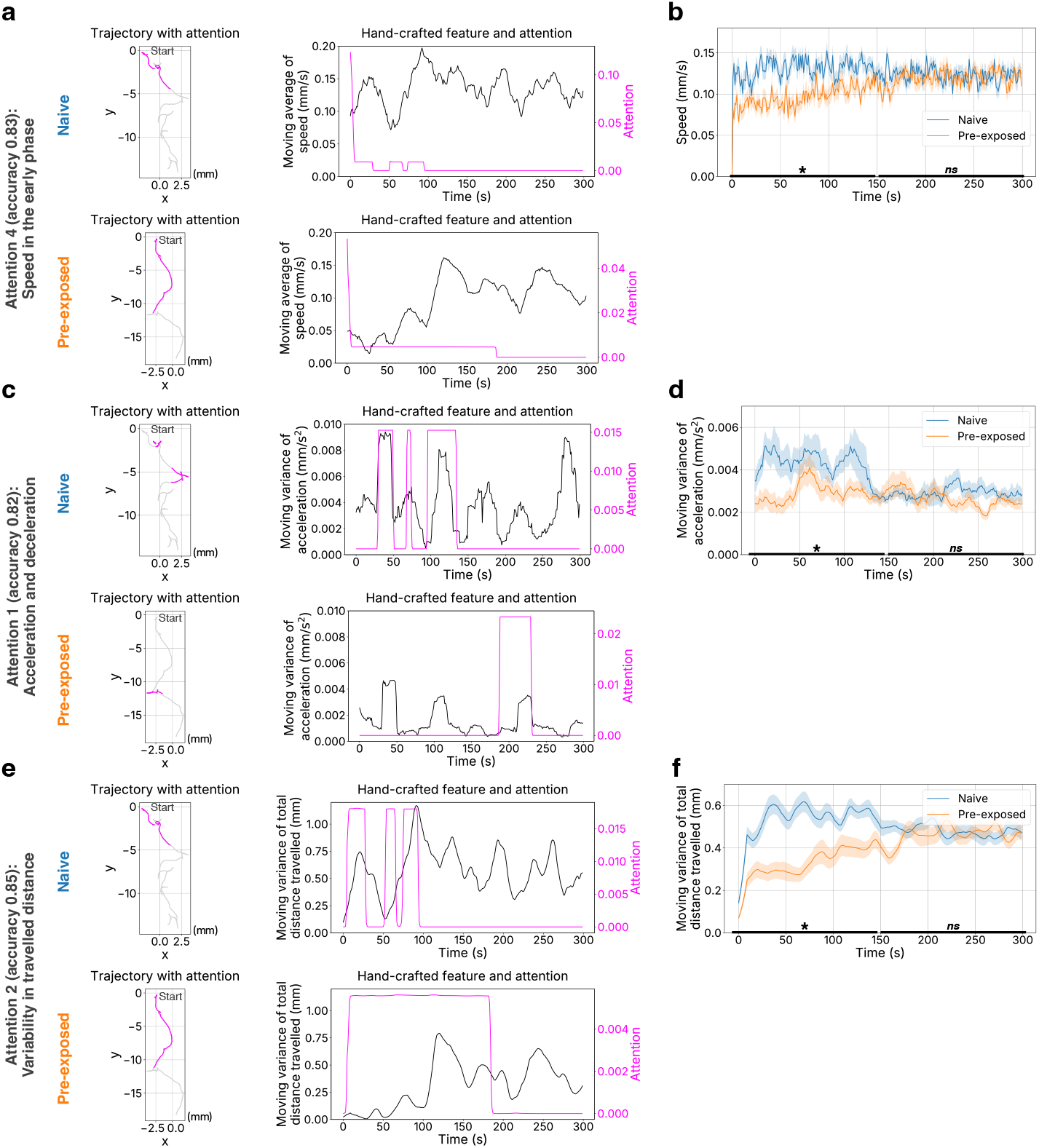
DiverScan rediscovers known features of worm odour-avoidance behaviour. Attention branches 4, 1, and 2 were associated with speed, acceleration variability, and variability in travelled distance, respectively. Post hoc tests recovered known signatures of odour learning: in the early phase, pre-exposed worms moved more slowly than naive worms and showed reduced fluctuations. **a,** Attention branch 4, paired with speed. Left: example trajectories with high-attention segments highlighted; right: the attention time series together with the associated hand-crafted feature. **b,** Time course of speed, compared between naive (*N* = 33) and pre-exposed (*N* = 33) trajectories. The mean values in the early and late phases (as defined in Fig. 3) were compared using two-sided Student’s *t*-tests with Holm correction (early phase, adjusted *p* = 2.8 10*^−^*^7^, Cohen’s *d* = 1.5, 95% CI [0.02, 0.05]; late phase, adjusted *p* = 0.29, *d* = 0.26, 95% CI [0.0036, 0.012]). Asterisks and *ns* denote *p* 0.05 and *p >* 0.05, respectively; solid lines and shaded regions indicate the mean and SEM, respectively. See Supplementary Table 4 for full statistics. **c,** Attention branch 1, paired with the moving variance of acceleration (moving window, 19 frames). **d,** Time course of the moving variance of acceleration (early phase, adjusted *p* = 0.013, *d* = 0.70, 95% CI [0.00038, 0.0022]; late phase, adjusted *p* = 0.34, *d* = 0.24, 95% CI [0.00033, 0.00094]; naive and pre-exposed, *N* = 33 each). **e,** Attention branch 2, paired with the moving variance of the total distance travelled (moving window, 19 frames). **f,** Time course of the moving variance of the total distance travelled (early phase, adjusted *p* = 1.5 *×* 10, *d* = 1.3, 95% CI [0.13, 0.28]; late phase, adjusted *p* = 0.49, *d* = 0.17, 95% CI [*−*0.04, 0.08]; *N* = 33 each).

**Fig. A4.**
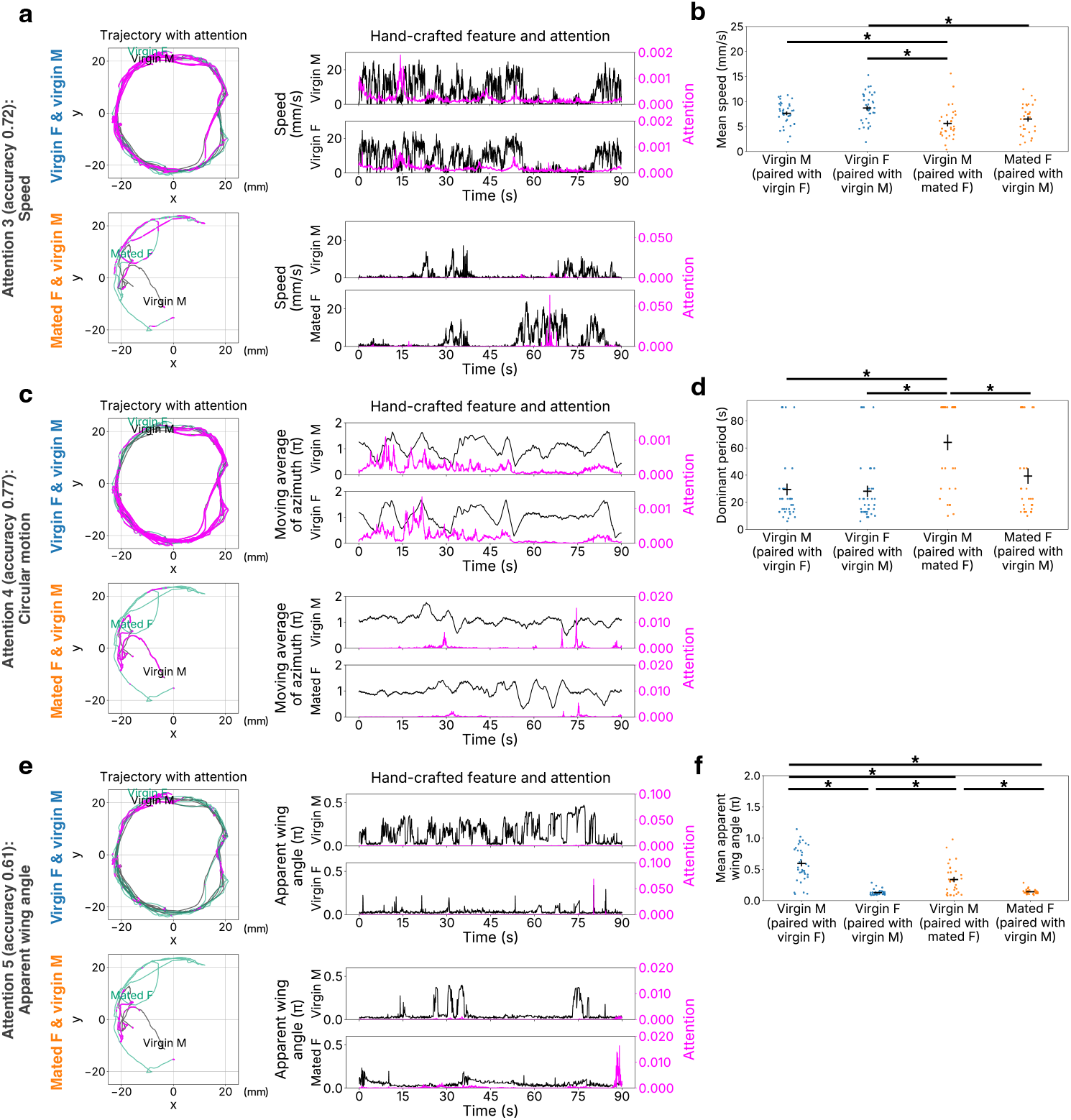
DiverScan rediscovers known features of fruit-fly courtship behaviour. Attention branches 3, 4, and 5 were associated with speed, the periodicity of movement, and the apparent wing angle, respectively. Post hoc tests recovered known courtship signatures: pairs with a virgin female showed higher speed and stronger movement periodicity, and the apparent wing angle differed with the partner’s mating status. **a,** Attention branch 3, paired with speed. Left: example trajectories with high-attention segments highlighted; right: the attention time series together with the associated hand-crafted feature. **b,** Mean speed across four conditions: virgin male with virgin female (VM–VF, 36 clips), virgin female with virgin male (VF–VM, 36 clips), virgin male with mated female (VM– MF, 31 clips), and mated female with virgin male (MF–VM, 31 clips). The six pairwise comparisons used two-sided Student’s or Welch’s *t*-tests (for equal or unequal group sizes, respectively) with Holm correction, listed in the order VM–VF vs. VF–VM, VM–VF vs. VM–MF, VM–VF vs. MF–VM, VF– VM vs. VM–MF, VF–VM vs. MF–VM, and VM–MF vs. MF–VM (adjusted *p* = 0.25, 0.020, 0.25, 8.2 10*^−^*^4^, 0.017, 0.26; Cohen’s *d* and 95% CIs in Supplementary Table 4). Asterisks denote adjusted *p* 0.05, and bars indicate the mean and SEM. **c,** Attention branch 4, paired with the distance from the initial location. Panels as in **a**. **d,** Dominant period of the distance-from-initial-location time series, estimated by fast Fourier transform. Conditions, tests, and comparison order as in **b** (adjusted *p* = 0.83, 5.7 10*^−^*^5^, 0.35, 2.6 10*^−^*^5^, 0.35, 0.013). **e,** Attention branch 5, paired with the apparent wing angle. Panels as in **a**. **f,** Mean apparent wing angle. Conditions, tests, and comparison order as in **b** (adjusted *p* = 4.8 *×* 10*^−^*^14^, 2.6 *×* 10*^−^*^4^, 8.9 *×*10*^−^*^11^, 1.2 *×* 10*^−^*^4^, 0.18, 1.2 *×* 10*^−^*^4^).

**Fig. A5.**
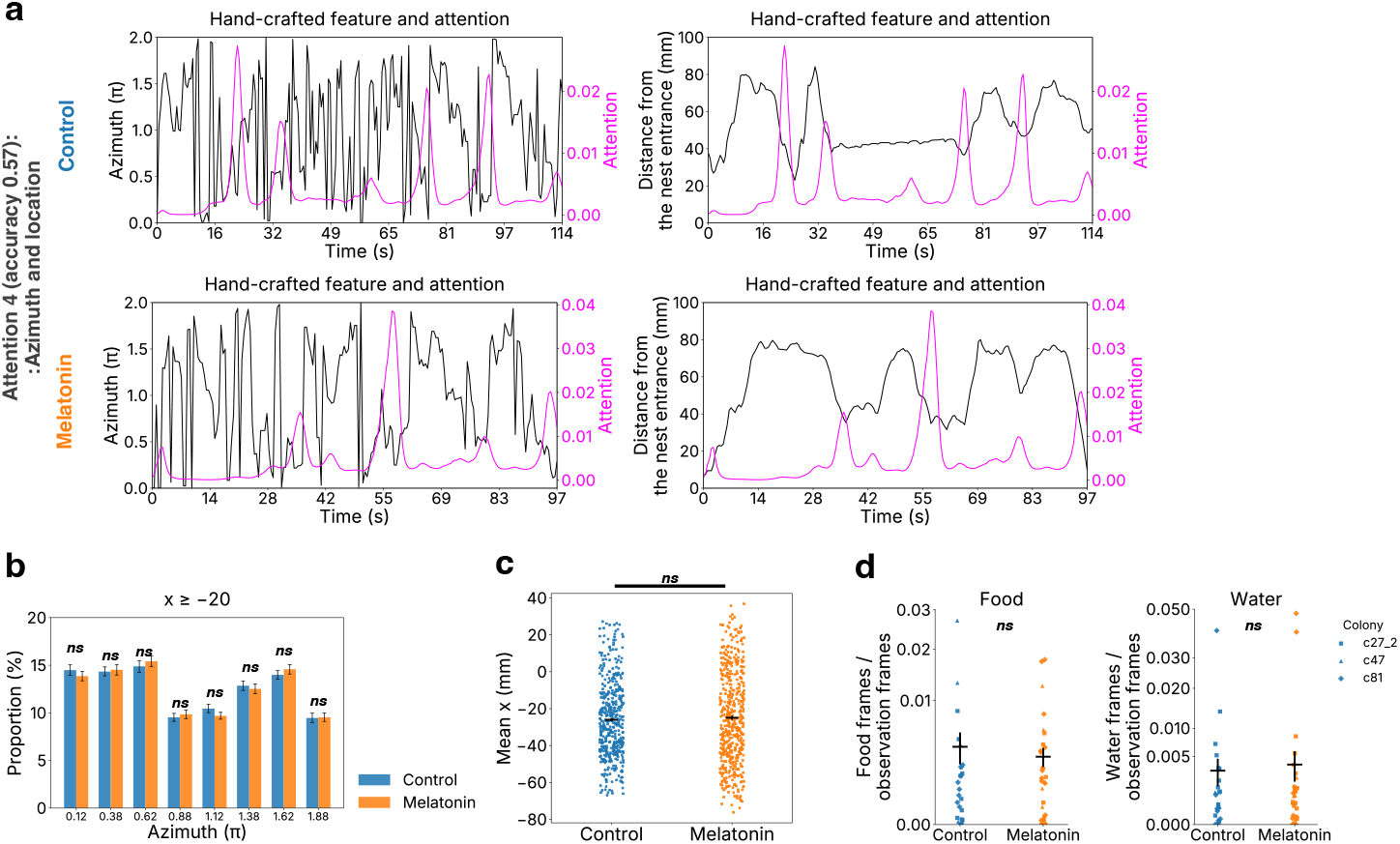
DiverScan points to a directional (azimuth) bias in isolated ants that conventional space-use metrics do not capture. Attention branch 4 was associated with the direction of movement (azimuth) and the distance from the nest entrance. Post hoc tests found no difference in mean *x* position or in time spent at the food or water supply, whereas the directional bias distinguished the conditions; see Supplementary Note for detailed discussion. **a,** Attention branch 4. Left: time series of the attention value and azimuth. Right: time series of the attention value and the distance from the nest entrance. Upper: *T ^′^* = 228 frames; lower: *T ^′^* = 195 frames (0.5 s per frame). **b,** Proportion of azimuths taken by ants in the region *x* 20 (the nest side), shown as a control for the *x <* 20 region analysed in Fig. 6d. Two-sided Welch’s *t*-tests with Holm correction (*N* = 445 control vs. 506 melatonin-treated sub-trips); effect sizes (Cohen’s *d*) and 95% CIs for all azimuth bins are given in Supplementary Table 4. *ns* denotes *p >* 0.05, and bars indicate the mean and SEM. **c,** Mean *x* position (two-sided Welch’s *t*-test, *p* = 0.45, Cohen’s *d* = 0.048, 95% CI [4.00, 1.79]; *N* = 445 control vs. 506 melatonin-treated sub-trips; bars indicate the mean and SEM). **d,** Per-day proportion of time spent at the food or water supply in isolated ants, with or without melatonin treatment (surviving individual*_√_*s <u>only</u>; control, *N* = 27; melatonin-treated, *N* = 32). The primary test was a linear mixed model, 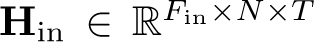) (food, *p* = 0.85; water, *p* = 0.91); a Wilcoxon rank-sum test was used as a sensitivity analysis because the residuals departed from normality (food, *p* = 1; water, *p* = 0.93). See Supplementary Table 4 for full statistics.

**Fig. A6.**
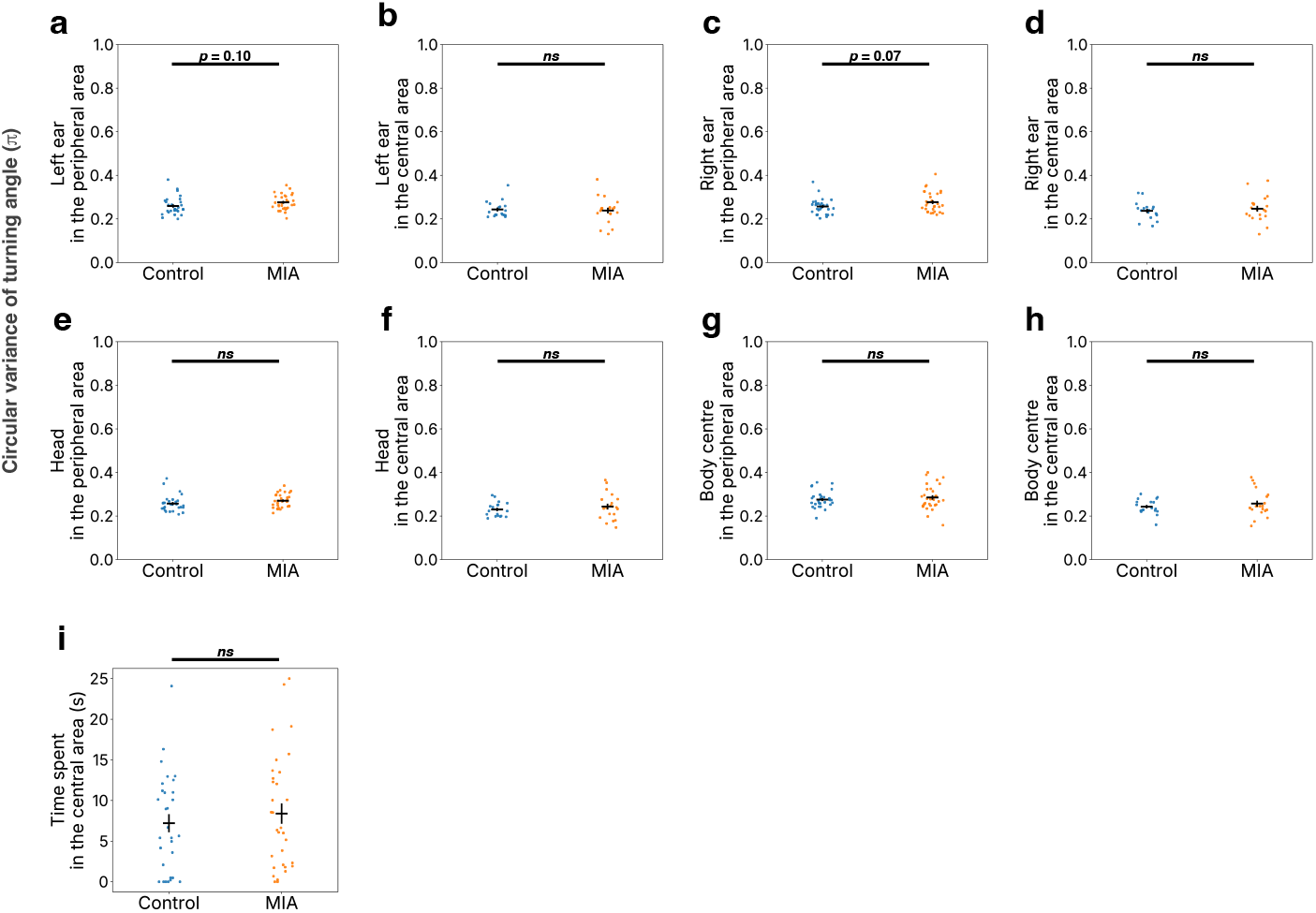
DiverScan points to a possible alteration in head-movement coordination in a rat model of schizophrenia. Panels **a**–**h** show the circular variance of the turning angle of a body part in the peripheral or the central area of the open field. All comparisons used two-sided Welch’s *t*-tests; *ns* denotes *p >* 0.05, and bars indicate the mean and SEM. See Supplementary Note for detailed discussion. **a,** Left-ear turning angle, peripheral area. The peripheral area was defined as the region more than 37.5 cm from the arena centre; clips in which a rat spent *<* 5 s there were excluded (*N* = 29 control and 30 MIA clips; *p* = 0.095, Cohen’s *d* = 0.44, 95% CI [0.037, 0.0031]). **b,** Left-ear turning angle, central area. The central area was defined as within 37.5 cm of the arena centre, with the same *<* 5 s exclusion (*N* = 18 control and 20 MIA clips; *p* = 0.76, *d* = 0.098, 95% CI [0.03, 0.04]). **c,** Right-ear turning angle, peripheral area (*p* = 0.071, *d* = 0.48, 95% CI [0.042, 0.0018]). **d,** Right-ear turning angle, central area (*p* = 0.56, *d* = 0.19, 95% CI [0.04, 0.02]). **e,** Head turning angle, peripheral area (*p* = 0.16, *d* = 0.37, 95% CI [0.03, 0.01]). **f,** Head turning angle, central area (*p* = 0.40, *d* = 0.27, 95% CI [0.04, 0.02]). **g,** Body-centre turning angle, peripheral area (*p* = 0.44, *d* = 0.20, 95% CI [0.03, 0.02]). **h,** Body-centre turning angle, central area (*p* = 0.38, *d* = 0.28, 95% CI [0.05, 0.02]). **i,** Time spent in the central area (*p* = 0.48, *d* = 0.18, 95% CI [4.5, 2.2]; *N* = 30 control and 32 MIA clips).

**Fig. A7.**
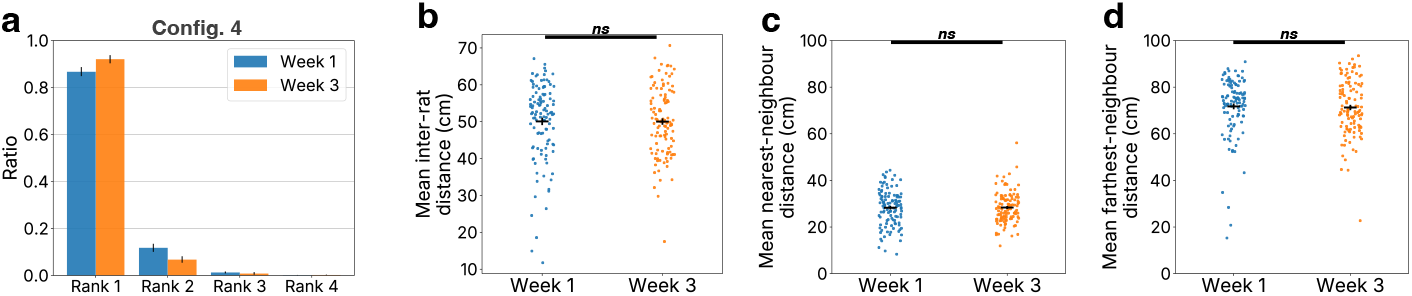
DiverScan captures group-configuration changes during social relationship formation in rats, undetected by simple distance metrics. **a**, Isolation ratio in configuration 4. For each clip, the time each rat spent isolated (i.e., the time points at which a single rat was separated from the other three) was computed in weeks 1 and 3 (112 clips each). Within each group, rats were ranked by this ratio, from rank 1 (longest isolation) to rank 4 (shortest); the consistently high ratio at rank 1 indicates that the same individual tended to be the isolated one. **b,** Mean inter-rat distance, compared between week 1 and week 3 with a two-sided Student’s *t*-test (*N* = 112 clips each; *p* = 0.97, Cohen’s *d* = 0.005, 95% CI [2.57, 2.68]). *ns* denotes *p >* 0.05, and bars indicate the mean and SEM. **c,** Mean nearest-neighbour distance (week 1 vs. week 3, *N* = 112 each; *p* = 0.92, *d* = 0.014, 95% CI [*−*2.00, 1.80]). **d,** Mean farthest-neighbour distance (week 1 vs. week 3, *N* = 112 each; *p* = 0.78, *d* = 0.037, 95% CI [*−*2.82, 3.74]).

**Fig. A8.**
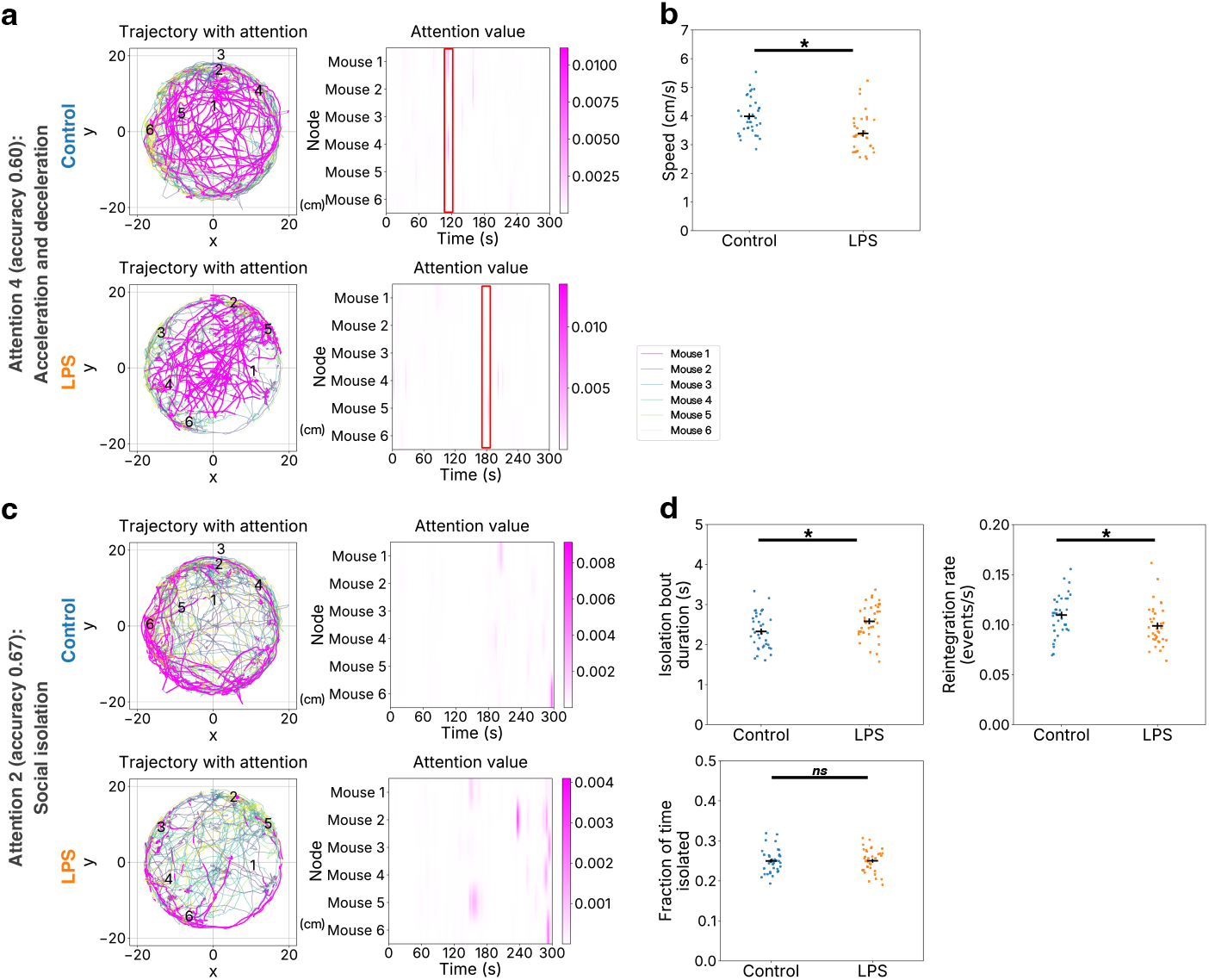
DiverScan points to altered movement dynamics in a mouse model of depression. Attention branch 4 was associated with acceleration and deceleration, and branch 2 with social isolation. Post hoc tests showed that depression-model mice had lower mean speed and longer isolation bouts with less frequent reintegration, while the total time isolated was unchanged. **a,** Attention branch 4 (acceleration and deceleration). Left: example trajectories with high-attention segments highlighted, where attention above the chance level 1*/*(*N T ^′^*) (*T ^′^* = 3,000 frames, *N* = 6 animals) is shown in magenta; right: heatmaps of attention values, with red rectangles marking the time windows shown in Fig. 6l. **b,** Mean speed, compared between control and depression-model (LPS) groups with a two-sided Student’s *t*-test (36 control vs. 36 LPS clips; *p* = 2.4 10*^−^*^4^, Cohen’s *d* = 0.91, 95% CI [0.29, 0.90]). The asterisk denotes *p* 0.05; horizontal and vertical bars indicate the mean and SEM, respectively. **c,** Attention branch 2 (social isolation), displayed as in **a** (without the chance-level threshold). **d,** Dynamics of social isolation. A mouse was scored as isolated at a given time point when its nearest-neighbour distance exceeded a threshold set to the 75th percentile of all pair-wise nearest-neighbour distances across the dataset; contiguous isolated time points form an isolation bout. Groups were compared with two-sided Student’s *t*-tests (36 control vs. 36 LPS clips); asterisks and *ns* denote *p* 0.05 and *p >* 0.05, respectively, and bars indicate the mean and SEM. Top left: mean isolation-bout duration, longer in the depression model (*p* = 0.018, Cohen’s *d* = 0.57, 95% CI [0.46, 0.04]). Top right: reintegration rate (transitions out of isolation per unit time), lower in the depression model (*p* = 0.027, *d* = 0.53, 95% CI [0.0012, 0.020]). Bottom left: fraction of time spent isolated, which did not differ between groups (*p* = 0.95, *d* = 0.016, 95% CI [*−*0.01, 0.01]).

**Fig. A9.**
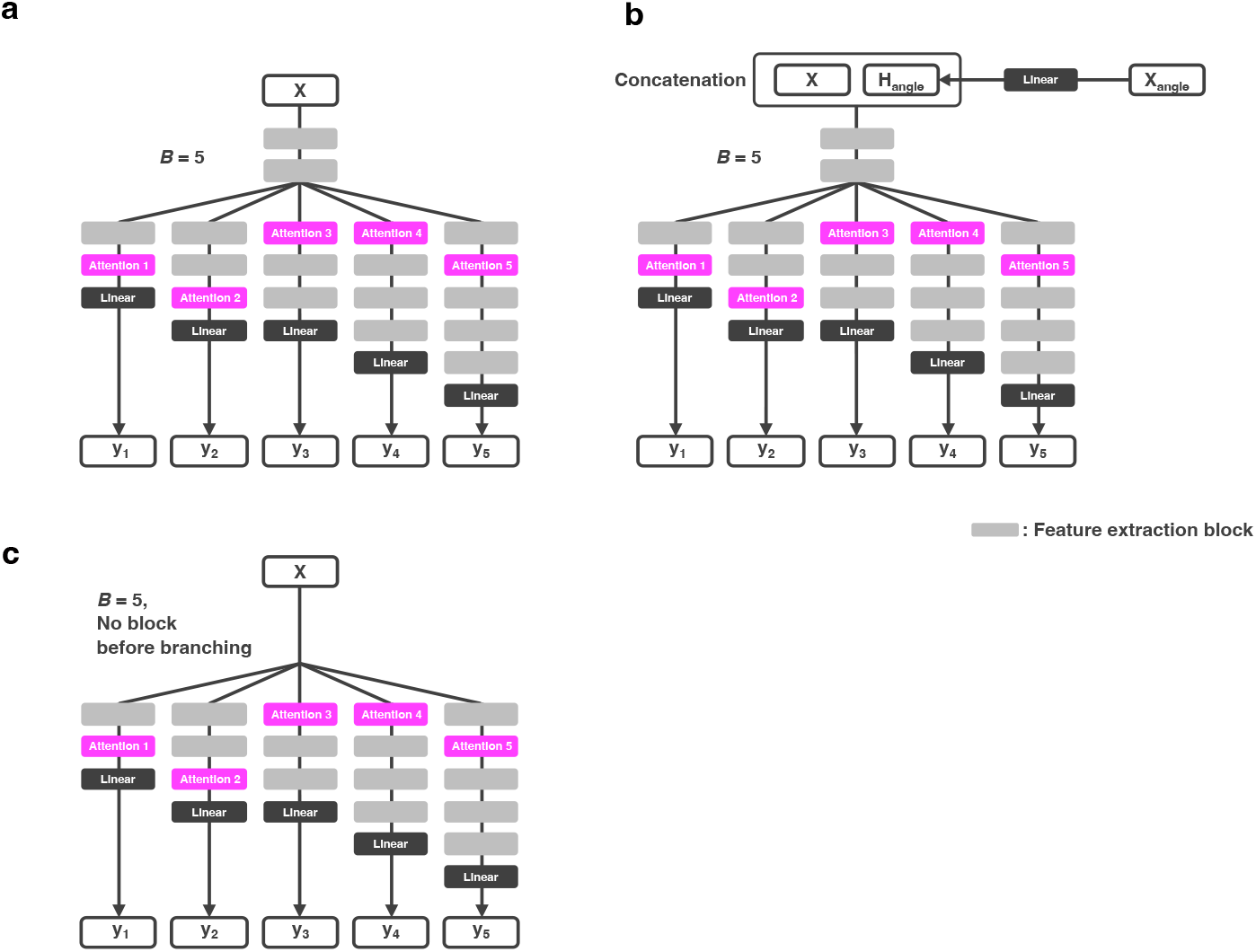
Example architectures of DiverScan-Net. **a**, A representative architecture that takes the input time series **X** directly, with feature-extraction blocks shared before the network branches. The network splits into *B* = 5 attention branches, each ending in a linear block that predicts the class (**y**_1_, *. . .,* **y**_5_); grey blocks are feature-extraction blocks, magenta blocks are attention blocks, and dark blocks are linear layers. Within each branch, the number of feature-extraction blocks and the position of the single attention block are chosen independently by the neural architecture search, so branches can extract different representations and attend at different depths (for example, attention branch 3 attends immediately after branching, whereas attention branch 2 attends after several feature-extraction blocks). **b,** An architecture that additionally uses the angles between body parts. The angle time series **X**_angle_ is passed through one linear block to give **H**_angle_, which is concatenated with **X** before entering the network. **c,** An architecture with no feature-extraction blocks before branching, so that the network splits into branches immediately after the input (the number of shared blocks is set to zero by the neural architecture search).

## Appendix B Supplementary note on the quantitative evaluation of DiverScan loss using synthetic signals

The purpose of this evaluation is to test whether DiverScan can correctly separate and detect distinct features on data with a known ground truth. Because the key features that distinguish conditions in real datasets are unknown, we constructed synthetic signals into which characteristic segments were deliberately inserted, and evaluated whether each attention branch isolates a different one of these segments. This controlled setting also lets us test how each component of DiverScan loss contributes to this separation, by removing or replacing the components one at a time.

### B.1 Dataset

Two types of synthetic signals were generated to evaluate DiverScan loss, as illustrated in Extended Data Fig. A1a:

- **Low-Mid-High condition**: Sine waves at 1 Hz (low frequency), at 6, 7, 8, and 9 Hz (mid frequency), and at 14 Hz (high frequency) were concatenated in random order, with each segment assigned a duration drawn uniformly from 5 to 10% of the signal length. Every signal contained at least one low-frequency, one mid-frequency, and one high-frequency segment.
- **Mid condition**: Sine waves at 6, 7, 8, and 9 Hz (mid frequency) were concatenated in random order with the same duration rule, but neither the 1 Hz nor the 14 Hz segments appeared.

The task is therefore to classify Low-Mid-High vs. Mid signals, for which the 1 Hz and 14 Hz segments are the only informative features; a successful model should attend to these two segments, ideally with one branch per feature. For both conditions, the sine-wave amplitude was set to 1.0 and Gaussian noise with standard deviation 0.1 was added on top of the wave. Each signal lasted 10 seconds at a sampling rate of 100 Hz, yielding *T* = 1,000 samples per signal. The 1 Hz and 14 Hz segments thus served as the distinguishing features of the Low-Mid-High condition. For each condition, 500 signals were generated (300 for training, 100 for validation, and 100 for testing), giving 1,000 signals in total. A short-time Fourier transform (STFT) was applied to each signal, and the magnitude of the resulting 6×1,001 spectrogram was used as the input to the model.

### B.2 Evaluation procedure

Identical network architectures and training settings were used across all compared approaches. The model consisted of a shared trunk of two 1D convolutional layers followed by *B* = 2 parallel branches, each composed of a single masked-attention block and a linear block. Models were trained for 30 epochs and the procedure was repeated for 5 random seeds. For each seed, the optimiser’s weight decay and the convolutional kernel sizes were tuned on the validation set with 15 Optuna trials; the architecture and loss configuration were held fixed across all seven approaches and all seeds. The constants appearing in the loss function were *C_L_* = 1, *C_S_* = 3, *C_w_* = 10, *T_L_*= 0.8 log 2 ≈ 0.555, *λ*_TV_ = *γ*_AD_ ≈ 3.96, *λ*_FC_ = 0.1, and *T*_AD_ = 0.8 (the threshold applied to the attention mass in the first or last 5% of the time axis).

We compared the following seven approaches, each defined by removing or replacing one component of DiverScan loss:

- **DiverScan**: DiverScan loss with all four components.
- **DiverScan w/o FC (Feature Coherence)**: the FC term is removed.
- **DiverScan w/o AD (Attention Distribution)**: the AD term is removed.
- **DiverScan w/o TV (Total Variation)**: the TV term is removed.
- **DiverScan w/o LP (Loss Penalty)**: the loss penalty is removed, i.e., *w_i_* = 1 for all *i*.
- **CE (Cross-Entropy) + Sim (Similarity)**: a loss defined as the simple sum of branch losses and branch similarities,

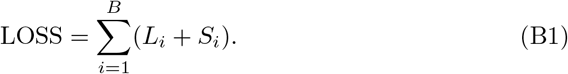

- **CE (Cross-Entropy)**: a loss defined as the sum of branch losses only,

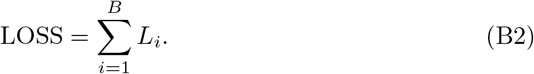

#### B.2.1 Performance during training

To check that each approach trains stably and to see how it balances classification against attention similarity, we monitored three quantities during training. For each approach, we monitored the validation values of the total loss LOSS, the averaged branch cross-entropy loss *L̄*, and the averaged branch similarity *S̄* across the 30 training epochs. The curves were summarised as the mean and standard error of the mean across the 5 random seeds and compared across the seven approaches.

#### B.2.2 Alignment of attention with the inserted features

We next asked whether, and how cleanly, the branches’ attention landed on the inserted low- and high-frequency segments. For each seed, the model checkpoint with the lowest validation loss across the 30 epochs was selected for test evaluation. To measure how well each branch’s attention aligned with the ground-truth features, we computed the Pearson correlation between each branch’s attention time series ***a****_i_*(*t*) and binary indicator sequences derived from the underlying frequency-of-segment time series ***f*** (*t*) ∈ {1, 6, 7, 8, 9, 14} Hz:

- mask_low_(*t*) = 1 ⇐⇒ *f* (*t*) = 1 Hz,
- mask_high_(*t*) = 1 ⇐⇒ *f* (*t*) = 14 Hz.

A correlation close to 1 indicates that the attention is well aligned with the corresponding frequency feature. Using a threshold *τ* = 0.5 on corr_low_ and corr_high_, each branch’s attention was classified into one of four states (the mid correlation was computed for reference only):

- **L** (low-frequency attention): corr_low_ *> τ* and corr_high_ ≤ *τ* ;
- **H** (high-frequency attention): corr_high_ *> τ* and corr_low_ ≤ *τ* ;
- **B** (both target attention): both correlations *> τ* ;
- **N** (neither): both correlations ≤ *τ* .

Because there are two branches (*B* = 2), the per-sample state was defined as the unordered pair {*s*_1_, *s*_2_} of the two branches’ labels, taking one of ten possible values. The pair {*L, H*} represents the ideal outcome, in which one branch attends exclusively to the low-frequency feature and the other exclusively to the high-frequency feature, so that the two distinguishing segments are cleanly separated into different branches. For each approach we report the fraction of correctly classified Low-Mid-High samples that fell into each state, averaged across the 5 seeds.

#### B.2.3 Purity, recall rate, and error rate

Reaching the {*L, H*} state tells us that each branch attends to the right feature overall, but not how precisely the highlighted time bins trace the target segments. To quantify this precision, we then focused on the subset of correctly classified Low-Mid-High samples whose two branches achieved the ideal state {*L, H*}, and quantified the purity, recall rate, and error rate of the time bins highlighted by the attention mechanism of each branch. A time bin was considered *highlighted* when its attention value exceeded 2*/T* here, twice the chance level 1*/T* obtained by a uniform attention over *T* = 1,001 STFT time bins. All three quantities were also re-computed after shrinking each target segment by 10% from each end, in order to test whether attention focused on the centre of a segment rather than on its boundary.

- **Purity**: a measure of whether an attention time series is dominated by a single frequency feature. We first counted the highlighted bins within the low-, mid-, and high-frequency segments separately, and then computed the ratio of the dominant (largest) count to the total number of highlighted bins. If the dominant count corresponded to the mid-frequency segments, the purity was set to 0.
- **Recall rate**: the fraction of low- or high-frequency time bins that were highlighted. For each branch, the metric was computed against both the low- and high-frequency binary masks.
- **Error rate**: the fraction of highlighted bins that fell outside the target frequency segments (e.g., highlighted mid-frequency bins).

### B.3 Results and discussion

Extended Data Fig. A1b shows the validation values of LOSS, the averaged branch cross-entropy loss *L̄*, and the averaged branch similarity *S̄* during training for all seven approaches. The total loss LOSS decreased under all approaches; the absolute scale differed across approaches because each formulation contains different terms. The averaged branch cross-entropy loss *L̄* also decreased under all approaches. The averaged branch similarity *S̄* remained relatively high under DiverScan, but this is expected rather than a failure to diversify: the informative segments (1 Hz and 14 Hz) can fall at similar positions in a signal, so even ideally separated branches show a similarity above the natural minimum, which corresponds to the segments lying at the two ends of the time series, for example 1 Hz first and 14 Hz last (Extended Data Fig. A1c). DiverScan drove the branch attentions apart only as far as the task required, without forcing them to be artificially dissimilar. Extended Data Fig. A1c shows the attention time series of the DiverScan model on a representative Low-Mid-High signal. The two branches successfully detected the inserted distinguishing segments and separated them: the low-branch focused on the 1 Hz (low-frequency) segments and the high-branch on the 14 Hz (high-frequency) segment, while neither branch attended to the mid-frequency segments. On a Mid signal (Extended Data Fig. A1d), which contains no 1 Hz or 14 Hz segments, both attention time series remained close to the chance level throughout.

Although classification accuracy on the test set was uniformly high across all approaches (≥ 96% for Low-Mid-High and essentially 100% for Mid in every seed), the branches differed markedly in how they distributed their attention. Extended Data Fig. A1e quantifies the per-sample branch state across correctly classified Low-Mid-High samples. The DiverScan approach reached the ideal {*L, H*} state for 100% of the samples, demonstrating that the two branches consistently and exclusively separated the low- and high-frequency features. Removing any single component of DiverScan loss degraded this separation. These results indicate that defining the loss as the product of *L_i_* and *S_i_*, rather than their sum, and adding the loss-penalty, total-variation, attention-distribution, and feature-coherence terms are each individually necessary to achieve robust branch separation in this synthetic dataset.

For samples in which both branches achieved the ideal {*L, H*} state, we further quantified the purity, recall rate and error rate of the highlighted time bins (Extended Data Fig. A1f–h). CE and CE + Sim were excluded from this analysis because they almost never reached the {*L, H*} state. The DiverScan approach achieved near-perfect purity (Extended Data Fig. A1f), together with high recall on the target bands (Extended Data Fig. A1g) and negligible error rates (Extended Data Fig. A1h). After shrinking each target segment by 10% from each end, the recall rose to ≥ 0.97, confirming that the small residual error of DiverScan loss was confined to segment boundaries, where the temporal extent of the convolutional receptive field unavoidably blurs the attention. Removing the TV term left purity largely unchanged but substantially lowered recall, reflecting attention curves that were sharply peaked rather than broadly covering the target segments and hence harder to interpret. Removing the FC, AD, or LP raised error rates, reflecting attention curves that frequently leaked into the mid-frequency segments.

Taken together, these results show that each component of DiverScan loss is necessary for interpretable feature extraction.

## Appendix C Supplementary notes for application of DiverScan in various experimental settings

### C.1 Activity in isolated ants

Social isolation is a common stressor in social species and is known to affect behaviour, physiological function, and ultimately lifespan. Behavioural alterations under social isolation have been documented across many animal systems. Using eusocial ants as a model, we previously focused on worker individuals and performed behavioural analyses based on automated tracking with individual identification using two-dimensional barcodes[27]. These studies showed that social isolation induces marked changes in space use compared with group-housed individuals: isolated individuals showed a pronounced reduction in nest occupancy and increased time spent in peripheral regions, particularly along the arena walls. These isolation-induced phenotypes are associated with an increase in oxidative-stress responses, which contribute to both behavioural abnormalities and shortened lifespan, and are alleviated by administration of the antioxidant melatonin.

However, previous analyses relied on partitioning the arena spatially and quantifying the time spent in predefined regions. Although such metrics capture large-scale spatial redistribution, they do not characterise movement trajectories in detail, such as directional dynamics or fine-scale behavioural structure. It therefore remained unclear whether isolation-induced behavioural changes involve altered directional preferences or biases towards specific environmental landmarks. Here we applied DiverScan to trajectory data from isolated ants, with and without melatonin treatment, to look for latent behavioural features not detectable with conventional space-use metrics.

We focused on attention branch 4, which attended to azimuth (direction of movement) and the distance from the nest entrance (Extended Data Fig. A5a). Although its classification accuracy was moderate (test-set accuracy 0.57; Supplementary Table 3), we highlight this branch because a directional signal is biologically interesting given the spatial layout of the assay; as elsewhere, we treated the attention as a starting point and verified its suggested features by post hoc analysis.

As described in the main text, in the region *x <* −20, the distribution of movement direction (azimuth) differed between conditions: control ants preferentially moved towards the food source, whereas melatonin-treated ants were biased towards the water source. By contrast, no clear condition-dependent difference was apparent in the region *x* ≥ −20, where the nest is located (Extended Data Fig. A5b). Consistent with this, the mean *x* position did not differ between isolated and melatonin-treated individuals (two-sided Welch’s *t*-test, *p* = 0.45, Cohen’s *d* = 0.048, 95% CI [−4.00, 1.79]; Extended Data Fig. A5c, Supplementary Table 4), indicating that the overall spatial distribution of individuals was comparable across conditions. Conventional space-use metrics likewise failed to detect a difference between treatments. In particular, we examined whether melatonin treatment affected the time that surviving isolated ants spent at the food or water supply—an analysis not included in our earlier study[28] and reported here for the first time. As the primary analysis, we used the per-day occupancy ratio, defined as the number of detected frames in each area divided by 172,800, and fitted a linear mixed model with colony identity as a random effect:

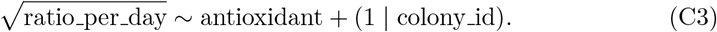

This revealed no significant effect of melatonin treatment for either area (food, LMM *p* = 0.85; water, LMM *p* = 0.91; Extended Data Fig. A5d, Supplementary Table 4). Because a Shapiro–Wilk test indicated that the LMM residuals departed from normality, we additionally performed Wilcoxon tests as a sensitivity analysis, both on the pooled data and separately for each colony; in every case, no significant difference was found for either food or water (Supplementary Table 4). By contrast, DiverScan pointed to a directional bias in movement as a distinguishing characteristic between conditions. These observations suggest that the behavioural difference between conditions is not necessarily reflected in spatial occupancy, but instead emerges in the directional structure of the movement trajectories.

This directional preference may relate to metabolic changes under isolation. Previous work showed that, although isolated individuals consume amounts of food comparable to those of group-housed ants, the amount of food transferred into the digestive tract is reduced[28], suggesting that ingested food is retained in the crop rather than being efficiently processed. In social insects, ingested food is first stored in the crop and later used for digestion or trophallaxis. The directional preference towards food or water sources identified here may therefore be linked to such metabolic alterations: one possibility is that changes in internal physiological state associated with reduced digestive efficiency are reflected in altered orientation towards specific resources, and that the shift in directional preference following melatonin treatment reflects a partial recovery of physiological homeostasis through reduced oxidative stress. Importantly, these differences were not detectable with conventional occupancy-time metrics and emerged only in the DiverScan analysis. Future work incorporating group-housed individuals will allow a more comprehensive understanding of how the social environment, interaction dynamics, and pharmacological intervention jointly shape behaviour and internal physiological state.

### C.2 Behaviour of a rat model of schizophrenia

Schizophrenia is a mental disorder shaped by both genetic and environmental factors[53]. Animal models have been widely used to investigate its genetic basis, environmental influences, and associated changes in brain function[54]. An important but often overlooked aspect of schizophrenia is abnormal motor behaviour, which can range from mild to severe and is closely linked to cognitive and emotional symptoms[29]. Difficulties with coordination and strength are common and can substantially reduce quality of life. However, quantifying coordinated movement across multiple body parts remains challenging, because diverse motor functions must be captured and interpreted simultaneously.

To address this challenge, we applied DiverScan’s multi mode to open-field recordings from two rat groups: a control group and a schizophrenia model induced by maternal immune activation (MIA). As input, DiverScan automatically constructed time-series graphs for 11 body parts (left ear, right ear, head, left shoulder, right shoulder, middle shoulder, body centre, left leg, right leg, middle leg, and tail). Nodes corresponded to individual body parts, edges were defined between all pairs of nodes, and edge features were given by the time series of inter-body-part distances. Because these body parts play fixed, asymmetric roles, we preserved the node ordering in the linear layer during training (i.e., we did not shuffle nodes; see Methods), so that DiverScan-Net could learn body-part-dependent latent representations. This setting lets DiverScan capture both the animal’s spatial position and fine body movements.

Among the attention branches, attention branch 2 achieved the highest classification performance (test-set accuracy 0.58; Supplementary Table 3). The accuracy was only moderate, so we treated the attention as a starting point and verified its suggested features by independent post hoc analysis. The attention-interpretation module suggested that this branch was associated with the distance from the centre of the open field, with particular emphasis on head-related body parts (the left ear, right ear, and head; Fig. 6f). The module also suggested that variability in inter-body-part distances was informative, possibly reflecting short-timescale movements. Collectively, these features could correspond to fine head-tilting motions. Post hoc analysis pointed to a tendency for MIA rats to show higher circular variance in ear turning angle in the peripheral area, although this did not reach significance (e.g., *p* = 0.095, Cohen’s *d* = 0.44, 95% CI [−0.037, 0.0031] and *p* = 0.071, Cohen’s *d* = 0.48, 95% CI [−0.042, 0.0018] for the two ears; Extended Data Fig. A6a–h, Supplementary Table 4). By contrast, we found no difference in the time spent in the centre of the open field (*p* = 0.48, Cohen’s *d* = 0.18, 95% CI [−4.5, 2.2]; Extended Data Fig. A6i, Supplementary Table 4), a commonly used index of anxiety-like behaviour, suggesting that the head-movement pattern is not simply a by-product of altered anxiety.

Together, these findings raise the possibility of impaired coordination of head movements in MIA rats—a feature not reported in the original study[30] but surfaced by DiverScan. Such fine motor changes would be consistent with aspects of schizophrenia pathophysiology: impaired motor coordination has been proposed as a potential endophenotype of schizophrenia, and a recent meta-analysis reported that youth with a family history of schizophrenia show delays in motor development, including impaired coordination, suggesting that motor skills may serve as early indicators[55]. Our results are therefore consistent with the possibility that fine motor-related markers could be informative for assessing schizophrenia risk and severity, a hypothesis that would need to be tested directly in future work.

